# Dose dependent impact of global *Cc2d1a* reduction on cognitive, social, and metabolic phenotypes

**DOI:** 10.64898/2026.10.03.756450

**Authors:** Abigail T. Heller, Aniket Bhattacharya, Pallavi Routray, Shanzeh Rauf, Vinitha Uppalapati, Emma Suzuki, Susanna Weiss, M. Chiara Manzini

**Author notes:** Corresponding author: M. Chiara Manzini, 89 French Street, Room 3274, Child Health Institute of New Jersey, New Brunswick, NJ 08901, USA.

## Abstract

**Background:** Autism spectrum disorder (ASD) and Intellectual disability (ID) share an increased male prevalence, but the biological mechanisms underlying this male bias remain poorly understood. Mice deficient for CC2D1A, a protein scaffold mutated in ASD/ID, have shown multiple male-specific deficits both in ASD-like behaviors and in the regulation of neuronal intracellular signaling pathways. However, previous research relied on conditional knockout (cKO) models due to early postnatal lethality of global *Cc2d1a* KO strains. Thus, the ability to define the developmental impacts of CC2D1A LOF in mice has been limited. While generating a *Cc2d1a* knock-in strain with V5-HA epitope tags, we found that homozygous animals (*Cc2d1a^VH/VH^*) resulted in a hypomorph line that can be used to study developmental loss of CC2D1A.

**Methods:** We assessed gross morphology, behavior, and intracellular signaling in male and female *Cc2d1a^VH/VH^* animals, where only ∼20% of CC2D1A is present. We then crossed this line with a null allele to obtain a more severe hypomorph (*Cc2d1a^-/VH^*) and explore phenotypic penetrance thresholds. In addition to the previous phenotypic characterization, we also assessed body composition, feeding, and molecular hallmarks of metabolic function in both sexes.

**Results:** We found that ∼80% CC2D1A reduction in *Cc2d1a^VH/VH^* mice did not recapitulate any of the behavioral or molecular disruptions seen in cKO models. In contrast, ∼90% CC2D1A reduction in *Cc2d1a^-/VH^* mice resulted in ASD/ID-like behavioral phenotypes in males, similar to those reported in cKO lines, with distinct intracellular signaling disruptions. Male and female *Cc2d1a^-/VH^* mice also showed subtle but significant adult weight deficits associated with distinct metabolic changes.

**Limitations:** These models still do not fully recapitulate the complete *CC2D1A* LOF found in patients. Due to the analysis of two independent strains over multiple metrics, these studies are not powered to compare statistical differences between sexes and data were analyzed in males and females independently.

**Conclusions:** We identified a strikingly narrow CC2D1A expression threshold for the emergence of ASD/ID-like behavioral deficits in males. In addition, we revealed a novel role for CC2D1A in adult weight gain. These mouse strains will be useful to better understand *CC2D1A* LOF phenotypes.

## Introduction

Biallelic loss-of-function (LOF) variants in *Coiled-coil and C2 domain-containing 1A* (*CC2D1A*) are a rare cause of fully penetrant Intellectual Disability (ID) with approximately 40% comorbidity with Autism Spectrum Disorder (ASD) (1–5). Affected individuals have minimal to no spoken language with seizures and sleep disturbances also reported (1–3, 6). ASD and ID are often comorbid with a complex genetic architecture where rare variants of high penetrance have been used to examine the molecular mechanisms of pathogenesis in mouse models (7–9). As both conditions have a markedly increased male prevalence with ASD having a 3.4:1 male:female diagnostic ratio and ID having a 1.7:1 diagnostic ratio (10, 11), genetic rodent models where sex-specific deficits arise, including *Cc2d1a* knock-outs (KO) have been studied to understand this sex bias (9, 12).

Previous work on multiple *Cc2d1a* conditional knockout (cKO) models successfully recapitulated core features of ASD and ID often revealing more severe behavioral and molecular deficits in males (13–17). Both removing *Cc2d1a* from glutamatergic neurons embryonically with *Emx1*-cre, and postnatally from the forebrain with *CaMKII*-cre show deficits in social behavior, aggression, hyperactivity, repetitive behaviors, avoidance, and multiple aspects of memory (13–19). When both male and female cKO animals were studied, cKO females had significantly fewer deficits (13, 17, 18). Interestingly, targeted postnatal removal of *Cc2d1a* from serotonergic neurons found treatment resistant anxiety and depressive behaviors but no sex differences (20, 21). Together, these studies emphasize an important role for CC2D1A in regulating affective, social, and cognitive function and a sex-specific effect that may originate from CC2D1A’s function in the forebrain.

CC2D1A is a multifunctional scaffolding protein that anchors to membranes via its C2 domain and coordinates protein–protein interactions through its *Drosophila melanogaster* 14 (DM14) domains (22, 23). DM14 domains are protein-binding domains that are unique to CC2D1A and its paralog CC2D1B. Studies in *Drosophila*, where there is only one ortholog lethal (2) giant discs (Lgd), and mouse models show that CC2D1A and CC2D1B act redundantly to preserve function (23–31). Double KO of *Cc2d1a* and *Cc2d1b* causes early embryonic lethality, but just one of four alleles can rescue this deficit (31). However, this functional redundancy is limited in neuronal contexts where *Cc2d1a* cKOs have social, affective, and cognitive deficits, while *Cc2d1b* KOs only show cognitive deficits (13–17). Additionally, *Cc2d1a-Cc2d1b* double heterozygotes act more similarly to the cKOs suggesting *Cc2d1a* plays larger role in neuronal regulation of behavior, possibly due to the fact CC2D1A is preferentially enriched in the post synaptic density compared to CC2D1B (16, 31, 32).

*In vivo* and *in vitro* studies in mouse models and human cell lines showed that CC2D1A plays a broad role in intracellular scaffolding proximal to membranes where it can promote binding of signaling proteins to regulate signal propagation (16, 17, 22, 30, 33–36). CC2D1A has been implicated in the regulation of various intracellular signaling pathways such as NF-κB, RAC1, PKA, and AKT (16, 22, 35–37). Studies have directly linked some of these signaling disruptions to behavioral deficits. Following AAV-mediated hippocampal *Cc2d1a* knock-down, activation of RAC1 and its downstream targets was increased and pharmacological inhibition of RAC1 rescued memory deficits (16). In forebrain-specific *Cc2d1a* cKO mice, male-specific behavioral impairments were associated with male-specific hyperactivation of cAMP scavenger, phosphodiesterase 4D (PDE4D), resulting in reduced PKA/CREB pathway signaling. Females remained unchanged, showing that CC2D1A regulates PKA/CREB signaling in sex-specific manner. Importantly, treatment with a PDE4D inhibitor rescued male-specific spatial memory deficits without affecting females (17).

While all these findings point to an important role for CC2D1A in controlling sex-specific signaling, their relevance to human disease is unclear due to the conditional *Cc2d1a* removal in mouse models. Multiple global *Cc2d1a* KO strains have been generated and pups are born at expected Mendelian ratios with normal in body size and brain morphology, but die within a few hours after birth due to breathing and swallowing deficits (14, 22, 28, 31, 34, 38). While cKO studies have been vital for understanding how intra-cellular signaling dysregulation is tied to behavioral deficits in *Cc2d1a* LOF, it is difficult to ascertain what deficits are driven by CC2D1A LOF or whether there may be secondary effects due to the restricted timing/region of conditional removal.

Here, we introduce and characterize two global and developmental *Cc2d1a* hypomorph mouse lines leading to up to 90% protein reduction and bypassing early postnatal lethality. The hypomorphs survive into adulthood with no gross sensory-motor deficits but show different sex- and dose-specific impacts of CC2D1A expression on cognition, intra-cellular signaling, and weight. We find that just 20% of CC2D1A expression is sufficient to support typical development and reduction to 10% reveals similar cognitive and social deficits to cKOs in males with different signaling disruptions. In addition, both males and females show mildly reduced growth after puberty that may originate from different metabolic changes. Overall, *Cc2d1a* severe hypomorphs will provide a novel model to understand the complex developmental impacts of *CC2D1A* LOF.

## Methods

### Animals

All mouse experiments were conducted in accordance with protocols approved by the Rutgers University Institutional Animal Care and Use Committee (IACUC; protocol #201900037) and followed National Institutes of Health guidelines. The *Cc2d1a* V5-HA knock-in mouse line (hereafter *Cc2d1a^VH/VH^*) was generated using CRISPR-mediated insertion of a 78 bp tag encoding both V5 and HA epitopes at the C-terminus of the *Cc2d1a* gene by the Rutgers Genome Editing Shared Resource. The tagged allele produces an amplicon length of 462 bp in length, compared to the 384 bp wild-type (WT) allele, and was generated on a C57BL/6J background. Correct insertion and orientation of the tag were verified by sequencing germline-transmitted alleles in first-generation offspring. The compound heterozygous line (*Cc2d1a^−/VH^*) was generated by crossing the *Cc2d1a^VH^* line with a *Cc2d1a* null line (*Cc2d1a^−/+^*) carrying an exon 11 gene trap that truncates the transcript and produces a non-functional protein (14). All behavioral cohorts were generated to include approximately equal numbers of mutant and WT littermates of both sexes.

For genotyping, 2 mm tail biopsies were digested overnight at 55°C in 200 µL SNET buffer (20 mM Tris-HCl, pH 8.0; 5 mM EDTA, pH 8.0; 400 mM NaCl; 1% SDS) containing 2 µL proteinase K (VWR, 76485-898). PCR reactions (50 µL) contained 5x GoTaq Flexi buffer (Promega, M890A), 100 µM dNTPs (New England Biolabs, N0447L), 50 µM forward and reverse primers (sequences available upon request), 1 mM MgCl₂ (Promega, A351B), 1.25 U GoTaq G2 Flexi DNA polymerase (Promega, M780A), and 1–2 µL template DNA. PCR amplification was performed using a 30-cycle touchdown protocol. Products were resolved on 1–2.5% agarose gels run at 90 V for 60 min (*Cc2d1a* null allele) or 90 min (*Cc2d1a^VH^*allele).

### Sensory-motor, Social and Cognitive Behavioral Assessment

All behavioral experiments were conducted on mice 10-14 weeks of age. Experimenters were masked to animal genotype throughout behavioral testing. Animals were habituated to the testing room and handled by the investigators for 7 days prior to the start of experiments. The *Cc2d1a^VH/VH^* and *Cc2d1a^−/VH^* cohorts were tested independently.

Initial sensory-motor assessment was performed using the SHIRPA protocol as described in (39). Open field testing was conducted under red light for 15 min in a 40 cm × 40 cm arena, as described in Seibenhener and Wooten (2015) (40). Three-chamber sociability and social novelty preference assays were performed under red light following the protocol of Moy et al. (2004) (41).

The Morris Water Maze (MWM) was conducted in a 120cm diameter metal tub filled with room temperature (20–25°C) water to a depth of 30cm. The test consisted of four phases. During Visible Platform Training, the platform was placed in the east quadrant just above the water level. Latency to reach the platform was recorded for drops at each starting quadrant with a maximum time of 90 s and latencies from each starting quadrant were averaged for each trial. Two trials were performed with an inter-trial interval of 1 hour. Hidden Platform Training consisted of three trials per day over three days for a total of nine trials. The platform remained in the east quadrant and was placed below the water level invisible to the animal. During the Probe Trial the platform was removed, and mice were released in the west quadrant and tracked for 60 s. Finally, during Reversal Training, the platform was placed in the west quadrant below the water level. Each mouse was placed into the east quadrant and latency to find the new platform location was measured up to 90 s for 4 trials with an inter-trial interval of at least 15 mins. For Visible, Hidden, and Reversal training the latency-to-platform were measured with a stopwatch by an experimenter masked to genotype, while time-in-quadrant for the Probe trial was recorded using ANY-Maze 6.36 (Stoelting).

### Feeding and Activity Analysis

Food intake was tested on single-housed mice by providing ∼15 g of standard chow (5058/PicoLab Diet 20) with *ad libitum* water and weighing remaining food after 24 h. Food preference testing compared intake after providing ∼15 g of standard chow and ∼15 g of banana-flavored BioServ Supreme Mini-Treats in separate dishes for 24 h.

Baseline locomotor and feeding behaviors were obtained using TSE Phenomaster metabolic cages. Mice were singly housed with *ad libitum* access to food and water beginning in the afternoon. Locomotor activity analysis began after 24 h of acclimation and was binned into 1 h intervals. Food intake was analyzed across light and dark cycles.

### Body Composition

Body composition was measured using an EchoMRI Body Composition Analyzer (E26-386-M), quantifying fat mass, lean mass, free water, and total water. Mice were weighed immediately prior to scanning.

### Signaling Analysis via Western Blot

Following euthanasia, hippocampi and cortices were rapidly dissected in cold 1x PBS, flash frozen in liquid nitrogen, and stored at −80°C. Tissues were homogenized using disposable pestles (VWR, 66001-104) in 400–600 µL of 1% SDS in PBS, supplemented with protease (Sigma-Aldrich, P8340) and phosphatase inhibitors (Sigma-Aldrich, P0044; 1:500). Homogenates were incubated on ice for 10 min and centrifuged at 15,000 rpm for 10 min at 4°C. Protein concentrations were determined using a BCA assay (G-Biosciences, 786-571). Samples were prepared in 4x Laemmli buffer (Bio-Rad, 1610747), and 30 µg protein was resolved on 8% or 4–12% Bis-Tris gels (Thermo Fisher Scientific, NW00082BOX, NW04125BOX). Proteins were transferred to nitrocellulose membranes (LI-COR, 926-31092) and blocked for 1 h at room temperature in either 3% BSA in TBS or 5% BSA in TBS-T (0.1% Tween-20). Membranes were incubated in primary antibody for 24-48 h at 4°C with rabbit anti-CC2D1A 1:1000 (Abcam, ab191472), rabbit anti-CC2D1B (Proteintech, 20774-1-AP), rabbit anti-phospho CREB (S133) (Cell Signaling Technology, 9198S), rabbit anti-CREB (Cell Signaling Technology, 9197L), mouse anti CaMKII-α (Thermo Scientific, MA1-048), and rabbit anti-phospho CaMKII (T286) (Cell Signaling Technology, 12716S), rabbit anti-phospho-PKA C (T197) (Cell Signaling Technology, 5661S), rabbit anti-PKA C α (Cell Signaling Technology, 5842S), mouse Anti-MAPK (ERK1/2) (Sigma Aldrich, M8159), rabbit p44/42 MAPK (ERK1/2) (Cell Signaling Technology, 4695), mouse anti-β Actin (Abcam, ab6276), mouse anti-GAPDH (Proteintech, 60004-1-IG). After primary antibody incubation membranes were washed thrice with TBS for 5 mins and incubated for 1 hour at room temperature with Alexa Fluor 680 Goat Anti-Mouse IgG (Invitrogen, A-21057) and Goat Anti-Rabbit IgG 800 (LICORbio, 926-3221). Membranes were then washed thrice with TBS-Tween 0.1% for 10mins. Blots were imaged using using LI-COR Odyssey CLx and analyzed with Licor ImageStudioLite software (V5.2.5, 2015). Target protein quantification was normalized to a GAPDH or β Actin loading control.

### qPCR Analysis

Brainstem tissue was dissected, flash frozen, and homogenized in 1mL TRIzol (Thermo Fisher Scientific, 15596026). RNA was extracted via chloroform separation and isopropanol precipitation, followed by ethanol washes. RNA pellet was dried and reconstituted in sterile water. cDNA was synthesized using iScript Reverse Transcription Supermix (Bio-Rad, 1708841) and diluted to 2ng/µL. qPCR was performed using PowerUp SYBR Green Master Mix (Thermo Fisher Scientific, A25776). Relative expression of *Cc2d1a* was calculated using the ΔΔCt method normalized to GAPDH (primers available upon request). For arcuate nucleus analysis, brains were sectioned (200–300 µm), and arcuate nuclei were micro dissected and stored in RNAlater (Thermo Scientific, AM7020). RNA was extracted using the Qiagen RNeasy Plus Kit (Qiagen, 74134) according to manufacturer protocol and qPCRs were performed as above. Expression of *Npy*, *Agrp, Pomc*, and *Cartpt* (*Cart*) was analyzed using published primers (42), and normalized to Actin and GAPDH.

### Hormone Analysis

Blood was collected mid-afternoon via submandibular bleed under isoflurane anesthesia. Blood was left at room temperature for 30 mins to clot and centrifuged at 1,500g for 10 mins. Serum supernatant was collected and stored at -20°C until analysis.

Serum hormone levels (TSH, prolactin, GH, FSH, ACTH) were measured using the MILLIPLEX Mouse Pituitary Panel (Millipore Sigma, MPTMAG-49K) and analyzed on a Luminex 200 (ThermoFisher, APX10031) system. Data were fit to a four-parameter logistic curve using Belysa software (v1.2) (Sigma Aldrich). Serum levels of corticosterone were determined using the DetectX® Corticosterone Multi-Format ELISA (Arbor Assays, K014-H1) 100 μl high sensitivity protocol according to manufacturer instructions.

### Statistical Analysis

Data are presented as mean ± SEM. Comparisons between two groups were performed using two-tailed Mann-Whitney U test or Welch’s t-test depending on data properties. Multiple group comparisons were analyzed using two-way ANOVA without an equal variance assumption (using the Geisser-Greenhouse assumption) followed by either a Šidák or Tukey’s post hoc test depending on data properties. Statistical significance was defined as p<0.05. All significance testing was performed using GraphPad Prism version 11.0.0 for Windows and Mac, Boston, Massachusetts USA, www.graphpad.com. Outliers in the qPCR and hormone analysis data sets are defined as any data point less than quartile 1 minus 1.5X interquartile range (IQR) and any data point above quartile 3 plus 1.5XIQR were removed. Outliers are marked in red in Data Master Spreadsheet (see data availability for link).

## Results

### Cc2d1a Hypomorph Line Development

Seeking to improve endogenous detection of CC2D1A, we generated a knock-in mouse strain where single V5 and HA epitope tags were inserted into the 3’ of the *Cc2d1a* coding region totaling a 78 base pair insertion (*Cc2d1a^VH^*, **Fig. 1A**). Unexpectedly, immunoblot analysis for the resulting protein revealed an approximate 80% reduction in total CC2D1A levels in *Cc2d1a^VH/VH^* homozygous animals (**Fig. 1B-C**) (Cortex. WT M: 0.855±0.172, n=4, *1a^VH/VH^* M: 0.181±0.021 n=6, *p*<0.0001, WT F: 1.112±0.084 n=5, *1a^VH/VH^* F: 0.236±0.011 n=7, *p*<0.0001, Hippocampus. WT M: 0.920±0.108 n=7, *1a^VH/VH^* M: 0.156 ± 0.018 n=7, *p*<0.0001, WT F: 1.062 ± 0.055 n=9, *1a^VH/VH^* F: 0.225 ± 0.047 n=11, *p*<0.0001). Analysis in early development additionally confirmed this reduction was present from birth and persistent into adulthood (**Fig. 1D**) (WT: 1.00±0.09 n=4, *1a^VH/VH^*: 0.16±0.05 n=7, *p*=0.0005). This protein reduction was present across multiple body regions tested in both nervous system and non-nervous system tissues (**Supp. Fig. 1**). Despite having only ∼20% of CC2D1A remaining, *Cc2d1a^VH/VH^* adult animals were viable and fertile. To our knowledge, *Cc2d1a^VH/VH^* mice represent the first model of global, developmental CC2D1A reduction that bypasses the early postnatal lethality observed in global *Cc2d1a* KO strains (14, 22, 28, 31, 34, 38).

**Figure 1.**
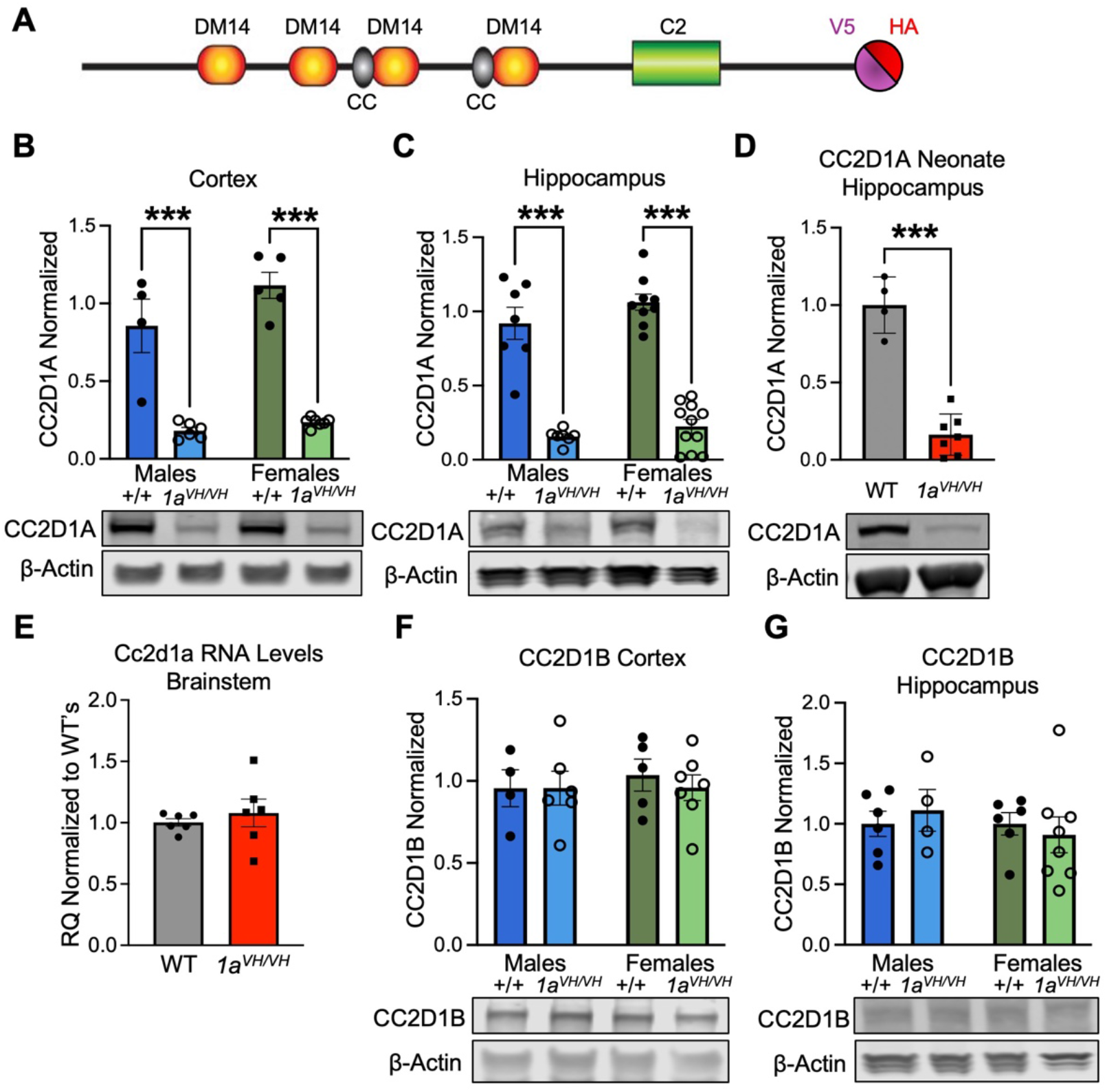
V5 and HA epitope tag insertion into C-terminus of CC2D1A causes protein degradation. **(A)** Diagram of V5 and HA epitope tags inserted into C-terminus of CC2D1A protein. Western blot shows *Cc2d1a^VH/VH^* mice display an ∼80% reduction in CC2D1A levels in adult **(B)** cortex and **(C)** hippocampus. **(D)** *Cc2d1a^VH/VH^*neonate hippocampi display the same significant reduction in CC2D1A. **(E)** qPCR revealed no significant changes in *Cc2d1a* transcripts (mRNA). Western blot shows no changes to total CC2D1B level in either adult **(F)** cortex or **(G)** hippocampus. CC2D1A levels are calculated by normalizing results to both male and female WT samples. Data are displayed as mean ± SEM. **(D, E)** Welch’s t test. **(B, C, F, G)** Two-way ANOVA with Fisher’s LSD. \*\*\**p*<0.001.

We wondered whether the tag insertion was disrupting mRNA stability. Quantitative PCR (qPCR) to examine *Cc2d1a* mRNA levels in the brain stem revealed no differences between genotypes, suggesting that the decrease occurs at the protein level, potentially due to destabilization caused by the epitope tags (**Fig. 1E**) (WT: 1.002±0.030 n=6, *1a^VH/VH^*: 1.079±0.113 n=6, *p*=0.309). Given the functional redundancy between CC2D1A and its homolog CC2D1B, we then asked whether compensatory upregulation of CC2D1B might occur (28–31). However, western blot analysis of adult cortical and hippocampal lysates showed that CC2D1B protein levels were comparable between *Cc2d1a^VH/VH^* mice and WT littermates, indicating no compensatory increase in expression (**Fig. 1F–G**) (Cortex. WT M: 0.956±0.113 n=4, *1a^VH/VH^* M: 0.956±0.103 n=6, *p*=0.999, WT F: 1.035±0.097 n=5, *1a^VH/VH^* F: 0.959±0.078 n=7, *p*=0.571, Hippocampus. WT M: 1±0.103 n=6, *1a^VH/VH^* M: 1.112±0.172 n=4, *p*=0.603, WT F: 1±0.093 n=6, *1a^VH/VH^* F: 0.909±0.148 n=8, *p*=0.614).

### Cc2d1a^VH/VH^ animals do not show physical or behavioral abnormalities

The unexpected viability of this mouse mutant allowed us to investigate how developmental reduction of *Cc2d1a* compares to previous conditional KO models. Initial analysis for body weight and gross sensory motor function using the SHIRPA protocol (39), including righting reflex, grip strength, nociception, and visual or auditory responses, showed no differences (**Fig. 2A**, **Supp. Table 1**) (Weight. WT M: 29.72±1.10g n=8, *1a^VH/VH^* M: 28.14±0.48g n=11, *p*=0.2966, WT F: 24.52±0.95g n=12, *1a^VH/VH^* F: 23.30±1.38g n=10, *p*=0.3816). We then asked how *Cc2d1a^VH/VH^*animals compare to *Cc2d1a* cKO models in their social and cognitive performance. In contrast to *Cc2d1a* cKO models targeting the forebrain or serotoninergic neurons, *Cc2d1a^VH/VH^* mice exhibited no alterations in total distance traveled (locomotion) or center avoidance in the open field test (**Supp Fig. 2 A-B**). (14–16, 20). Three-chamber sociability testing displayed no differences in sociability preference between an empty cup or stranger mouse (**Fig. 2B-C, Supp. Table 2**). In the Morris Water Maze (MWM) test for spatial memory, *Cc2d1a^VH/VH^* male and female mice showed no time to platform impairments during memory acquisition training (**Fig. 2D-E, Supp. Table 3**). *Cc2d1a^VH/VH^*mice did not display time in escape quadrant differences during the spatial memory retention portion of the MWM test (Probe trial) and had no significant delays in finding the platform during the cognitive flexibility trials (Reversal trial) (**Fig. 2F-H, Supp. Table 3**). Overall, none of the locomotor activity, sociability, and cognitive deficits identified in previous cKO studies were present in *Cc2d1a^VH/VH^*animals (13–17, 20).

**Figure 2.**
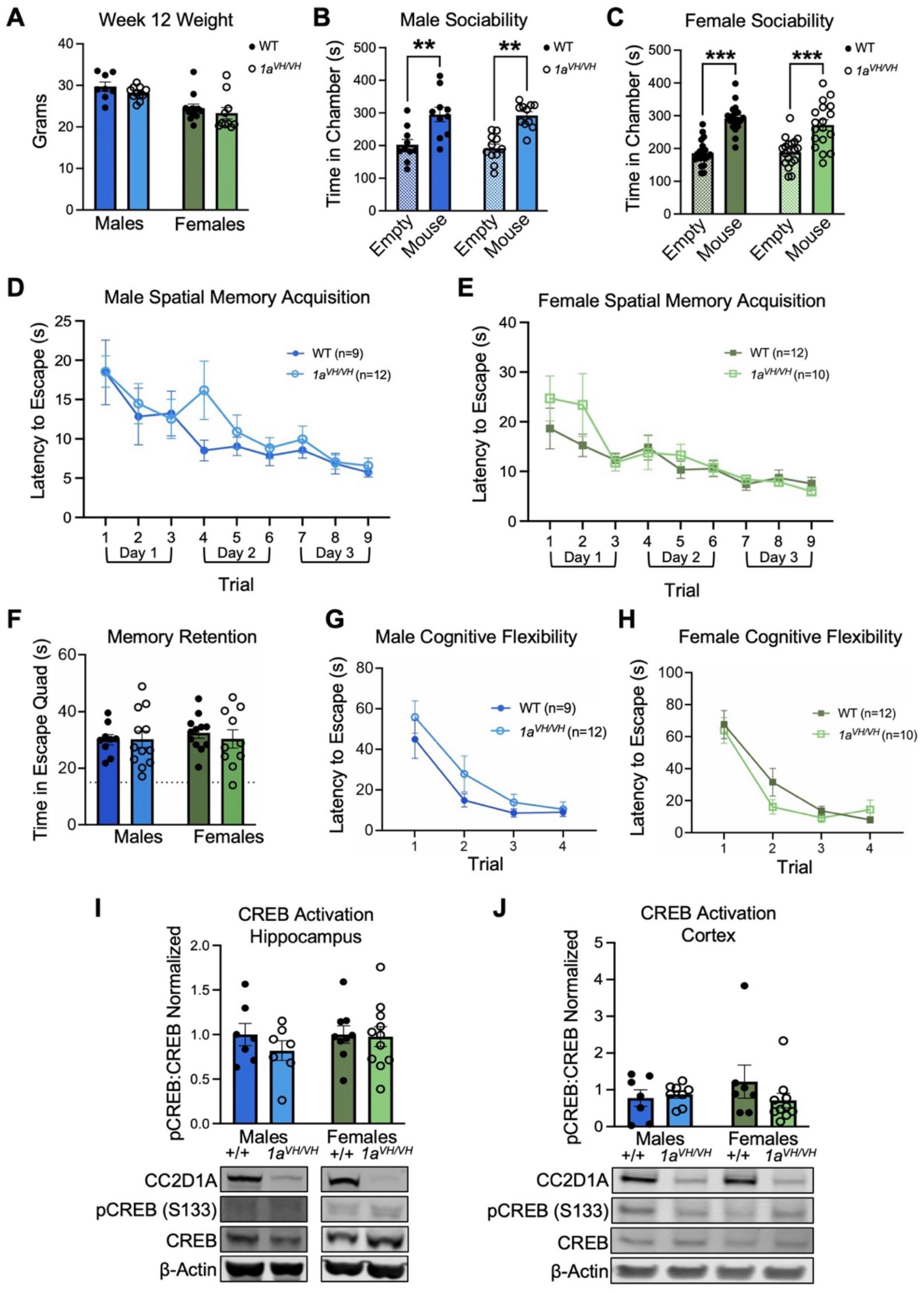
Cc2d1a^VH/VH^ mice do not display behavioral or baseline CREB deficits. **(A)** *Cc2d1a^VH/VH^*mice are comparable weight to WT littermates. Neither *Cc2d1a^VH/VH^*males **(B)** or females **(C)** display decreased sociability in 3-chamber social testing. Neither *Cc2d1a^VH/VH^* males **(D)** or females **(E)** display deficits finding the platform during MWM hidden platform training. Nor were there deficits during MWM probe **(F)** or reversal trials in males **(G)** or females **(H)**. Western blot revealed no differences in CREB activation in either adult hippocampus **(I)** or cortex **(J)**. Data are displayed as mean ± SEM. **(A-C, F, I-J)** Two-way ANOVA with Fisher’s LSD. **(D-E, G-H)** Two-way ANOVA with Šidák multiple comparisons test. \**p*<0.05, \*\*\**p*<0.001. WT, wild type; pCREB, S133 phosphorylated CREB.

We next checked if developmental CC2D1A reduction in the *Cc2d1a^VH/VH^* mice had any effect on the signaling pathways involved in cKO behavioral deficits. CREB signaling, was found to drive deficits in *Cc2d1a^CaMKII-Cre^* cKO models (17). However, we assessed baseline CREB activation via western blot by the ratio of CREB phosphorylation at serine 133 (S133) to total CREB. We found that it was unchanged in both the hippocampus and cortex of *Cc2d1a^VH/VH^* mice (**Fig. 2I–J**) (Hippocampus. WT M: 1.005±0.125 n=7, *1a^VH/VH^* M: 0.832±0.113 n=7, *p*=0.333, WT F: 0.996±0.099 n=9, *1a^VH/VH^* F: 0.973±0.111 n=11, *p*=0.876, Cortex. WT M: 0.776±0.223 n=7, *1a^VH/VH^* M: 0.875±0.103 n=8, *p*=0.792, WT F: 1.224±0.451 n=7, *1a^VH/VH^* F: 0.709±0.196 n=10, *p*=0.160). Together these data show that mice are remarkably resilient to CC2D1A reduction and that just 20% of the protein appears sufficient to prevent both behavioral and intracellular disruptions.

### CC2D1A expression reduction to 10% reveals behavioral deficits in males

The stark contrast between perinatal lethality with *Cc2d1a* loss in global KO strains and the results from *Cc2d1a^VH/VH^* animals led us to further reduce CC2D1A levels by crossing the *Cc2d1a^VH^* line (*Cc2d1a^+/VH^*) with heterozygous animals for a null allele (*Cc2d1a^+/-^*) generating compound heterozygotes (*Cc2d1a^-/VH^*) (**Fig. 3A**). This resulted in a near 90% reduction in CC2D1A expression (**Fig. 3B**) (WT M: 0.999±0.073 n=8, *1a^-/VH^*M: 0.133±0.023 n=10, *p*<0.0001, WT F: 1.001±0.101 n=6, *1a^-/VH^*F: 0.125±0.007 n=6, *p*<0.0001). As in *Cc2d1a^VH/VH^* mice, CC2D1B protein levels were unchanged, indicating no compensatory upregulation (**Fig. 3C**) (WT M: 1.112±0.109 n=6, *1a^-/VH^* M: 1.157±0.193 n=7, *p*=0.819, WT F: 0.888±0.136 n=6, *1a^-/VH^* F: 1.119±0.059 n=6, *p*=0.267).

**Figure 3.**
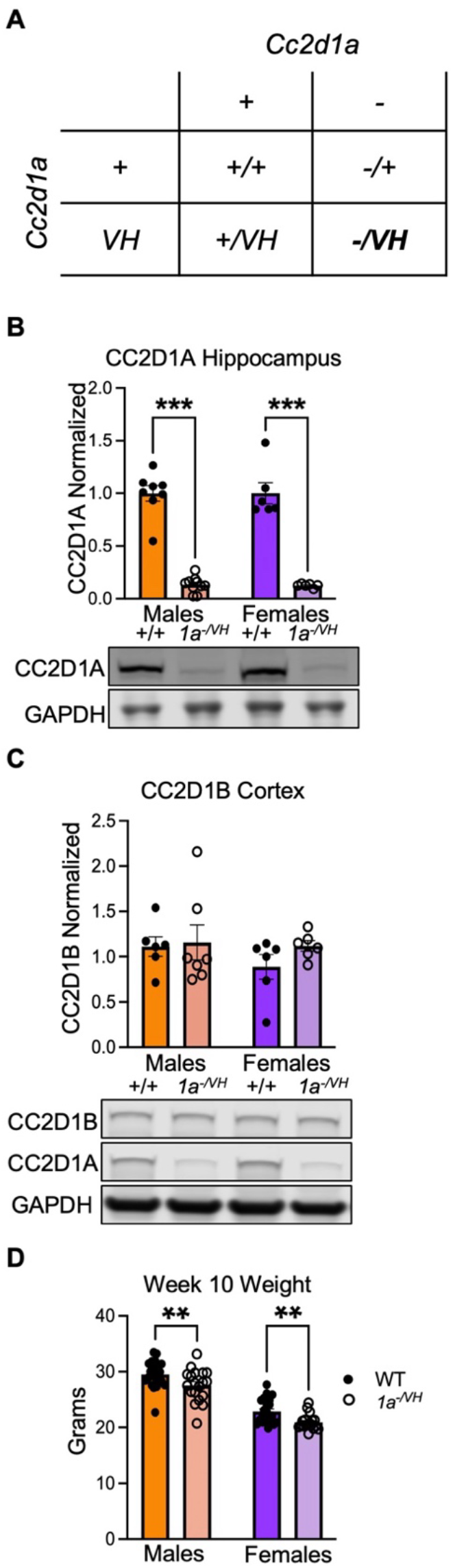
Generation of a more severe, 90%, *Cc2d1a* hypomorph. **(A)** Punnett square of *Cc2d1a^-/VH^* compound heterozygous breeding scheme. **(B)** Western blot of adult hippocampi reveal a near 90% reduction in total CC2D1A in *Cc2d1a^-/VH^* mice. **(C)** total CC2D1B levels remain unchanged in *Cc2d1a^-/VH^*adult cortex. **(D)** *Cc2d1a^-/VH^* mice are significantly underweight compared to WT littermates at 10 weeks old. Data are displayed as mean ± SEM. **(B-D)** Two-way ANOVA with Fisher’s LSD. \*\**p*<0.01,\*\*\**p*<0.001. WT, wild type; P, postnatal day.

SHIRPA analysis showed no gross sensorimotor deficits (**Supp. Table 4**). However, adult *Cc2d1a^-/VH^* mice displayed reduced body weight compared to WT littermates (**Fig. 3D**) (WT M: 29.51±0.54g n=21, *1a^-/VH^* M: 27.55±0.66g n=19, *p*=0.0085, WT F: 22.89±0.46g n=22, *1a^-/VH^* F: 20.95±0.33g n=18, *p*=0.0098). We found no differences in locomotion (total distance) or avoidance (time in center) behavior in the open field (**Supp. Fig. 3A–B**). However, in the three-chamber assay, male *Cc2d1a^-/VH^* mice showed a loss of sociability preference between the empty cup and the novel mouse (**Fig. 4A–B, Supp. Table 5**), similar to deficits observed in CaMKII-Cre and Emx1-Cre cKOs (14, 15). This was accompanied by *Cc2d1a^-/VH^* impairments in social novelty preference with a loss of discrimination between a familiar and novel mouse in males (**Supp. Fig. 3C–D, Supp. Table 5**). In the MWM, male *Cc2d1a^-/VH^* mice showed a trend for a delay in spatial memory acquisition during early hidden platform training (**Fig. 4C–D, Supp Table 6**). Focusing in on the last trial of each day shows this delay to be significant on the final trials of day one (trial 3) and two (trial 6), however this delay was resolved with additional training by day three as seen in cKOs (**Fig. 4E–F, Supp Table 7**). No differences were observed in probe trial performance or reversal learning (**Fig. 4G–I, Supp. Table 6**) (WT M: 26.62±1.61s n=15, *1a^-/VH^* M: 24.53±2.00s n=17, *p*=0.411, WT F: 25.86±1.74s n=13, *1a^-/VH^* F: 24.51±1.99s n=13, *p*=0.629). This pattern closely resembles *Cc2d1a^CaMKII-Cre^* cKO males, which had deficits in memory acquisition during early training that were resolved with further training (14, 17). Together, these findings identify a critical threshold for CC2D1A expression needed to impair sociability and cognition in males.

**Figure 4.**
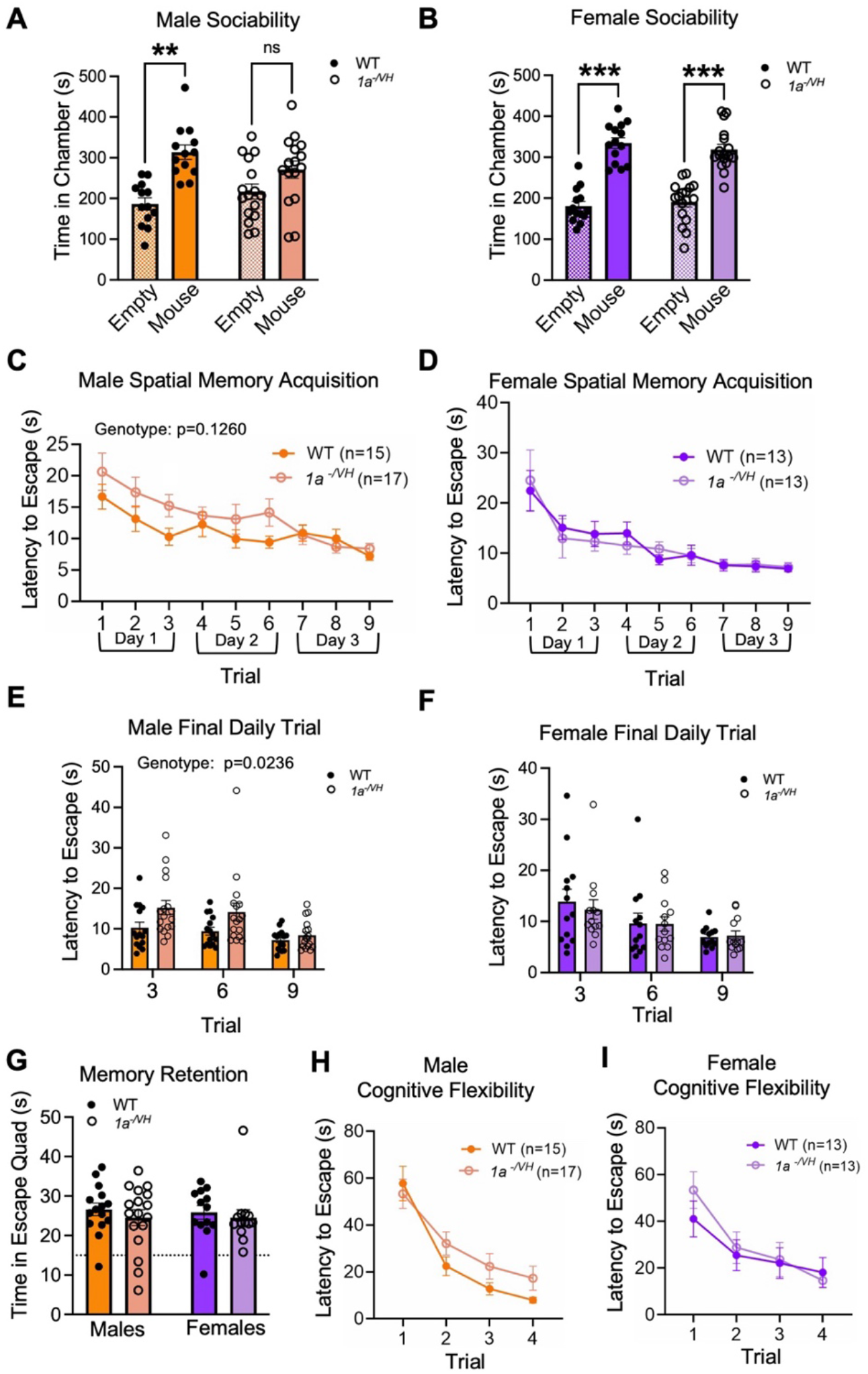
Male *Cc2d1a^-/VH^*mice display social and cognitive deficits. *Cc2d1a^-/VH^* males **(A)**, but not females **(B)**, have decreased sociability preference in 3-chamber sociability testing. **(C)** *Cc2d1a^-/VH^* males have significantly increased time to platform on trial 2 of hidden platform training **(D)**. While *Cc2d1a^-/VH^* females have no such spatial memory deficits. Examining the last trial of each day shows *Cc2d1a^-/VH^* males **(E)** but not females **(F)** have significantly increased time to platform trials 3 and 6. **(G)** There were no differences in time in escape platform between genotype or sexes, during the MWM probe trial. There were no significant delays in time to platform during cognitive flexibility reversal training in either males **(H)** or females **(I)**. Data are displayed as mean ± SEM. **(A-B, G)** Two-way ANOVA with Fisher’s LSD. **(C-F, H-I)** Two-way ANOVA with Šidák multiple comparisons test. **p<0.01, \*\*\**p*<0.001. WT, wild type.

### CC2D1A expression reduction to 10% leads to CaMKII disruption in males

Behavioral deficits in *Cc2d1a^CaMKII-Cre^* cKO males have been linked to hyperactivation of the cAMP-degrading enzyme PDE4D, resulting in reduced cAMP levels, decreased PKA activity, and diminished CREB activation (17). Given the behavioral similarities between *Cc2d1a^-/VH^*mice and *Cc2d1a^CaMKII-Cre^* cKOs, we examined whether similar signaling disruptions were present. Surprisingly, western blot analysis revealed baseline CREB phosphorylation at S133 was unchanged in both adult cortex and hippocampus (**Fig. 5A–B**) (Hippocampus. WT M: 0.839±0.135 n=8, *1a^-/VH^* M: 0.805±0.128 n=10, *p*=0.8889, WT F: 1.215±0.254 n=6, *1a^-/VH^* F: 1.188±0.279 n=6, *p*=0.9265, Cortex. WT M: 1.099±0.179 n=10, *1a^-/VH^* M: 1.292±0.237 n=13, *p*=0.5350, WT F: 0.890±0.113 n=9, *1a^-/VH^* F: 1.511±0.317 n=9, *p*=0.0798). Similarly, we found no differences in the activation of PKA at phosphorylation site T197 (**Supp. Fig. 4A-B**).

**Figure 5.**
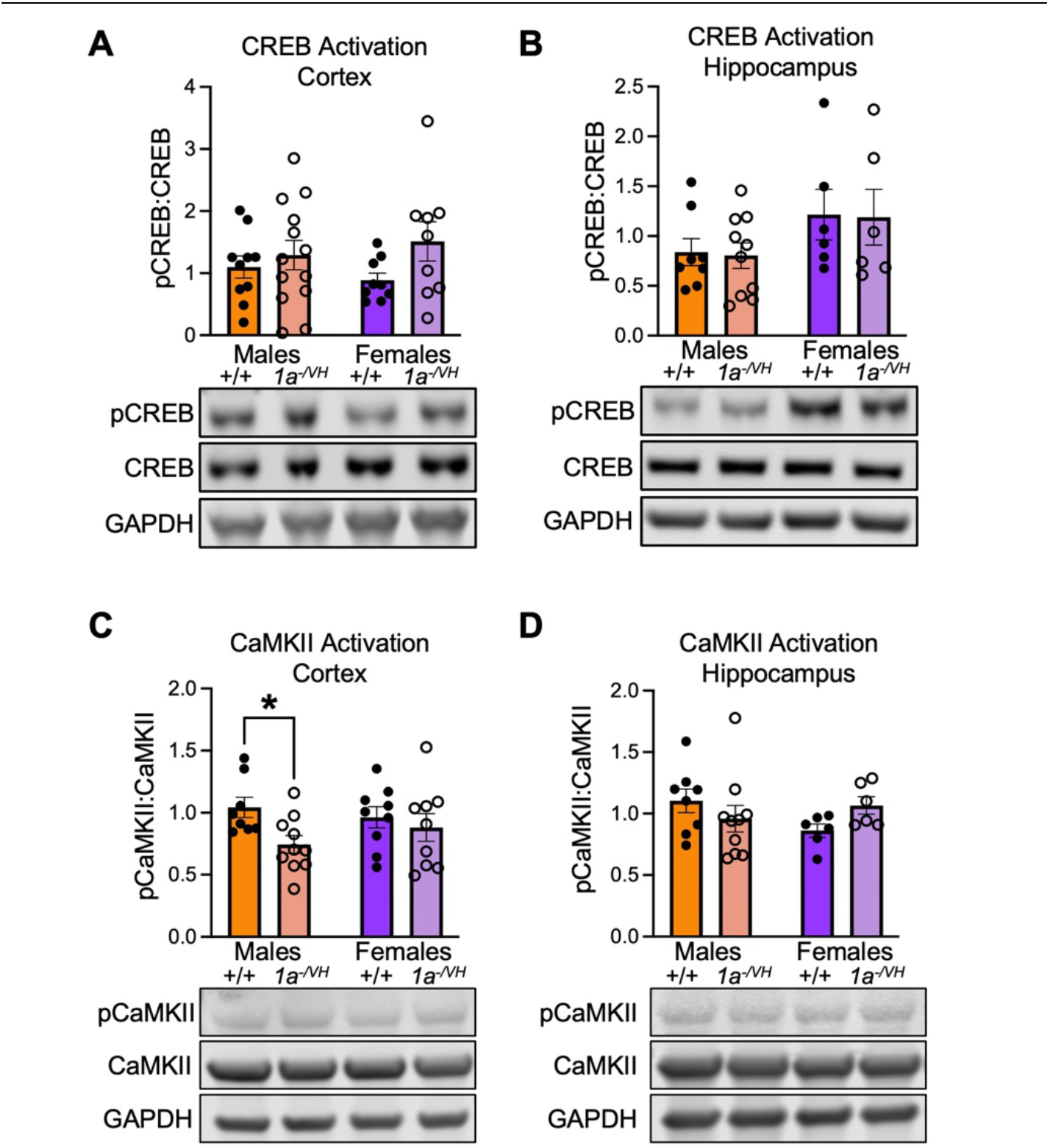
Male *Cc2d1a^-/VH^* mice display intracellular signaling deficits. Western blot adult cortex **(A)** and hippocampus **(B)** show no baseline CREB activation deficits. Western blot shows *Cc2d1a^-/VH^* males have decreased CaMKII activation in the adult cortex **(C)**, but not hippocampus **(D)**. Data are displayed as mean ± SEM. **(A-D)** Two-way ANOVA with Fisher’s LSD. pCREB, S133 phosphorylated CREB; pCaMKII, T286 phosphorylated CaMKII.

Since CC2D1A can regulate a variety of signaling pathways by altering membrane receptor trafficking and activation, we then investigated other signaling pathways involved in cognitive performance (30, 33, 34, 43–46). There was no change in the activation of ERK1/2 at phosphorylation sites T202/Y204 in either cortical or hippocampal adult lysates (**Supp. Fig. 4C-D**). Interestingly, analysis of CaMKII phosphorylation at T286 revealed a reduction in CaMKII activation in the cortex, but not the hippocampus, of male *Cc2d1a^-/VH^* mice (**Fig. 5C–D**) (Hippocampus. WT M: 1.103±0.096 n=8, *1a^-/VH^*M: 0.959±0.107 n=10, *p*=0.2558, WT F: 0.862±0.054 n=6, *1a^-/VH^* F: 1.066±0.071 n=6, *p*=0.1891, Cortex. WT M: 1.042±0.082 n=8, *1a^-/VH^* M: 0.744±0.071 n=10, *p*=0.0240, WT F: 0.963±0.084 n=9, *1a^-/VH^* F: 0.880±0.112 n=9, *p*=0.5141). Over all these results suggest that a ∼90% global reduction in CC2D1A recapitulates key behavioral phenotypes of *Cc2d1a^CaMKII-Cre^* cKOs but without baseline changes in PKA/CREB and a reduction in cortical CaMKII activation in males.

### Cc2d1a^-/VH^ mice have adult-onset weight deficits independent of feeding

We next investigated the unexpected weight deficit observed in *Cc2d1a^-/VH^*mice. To determine when this phenotype emerges, body weight was measured across multiple developmental time points starting at birth for both males and females. We found that *Cc2d1a^-/VH^* mice were born at weights comparable to WT littermates and maintained similar body weights in early development, but male and female *Cc2d1a^-/VH^* weight deficits emerged by 8 and 10 weeks of age, respectively (**Fig. 6A, Supp. Table 8**). While statistically significant these weight deficits are quite subtle with *Cc2d1a^-/VH^* males weighing 6.65%±2.23 less, and *Cc2d1a^-/VH^* females weighing 8.45%±1.43 less than WT littermates at 10 weeks old.

**Figure 6.**
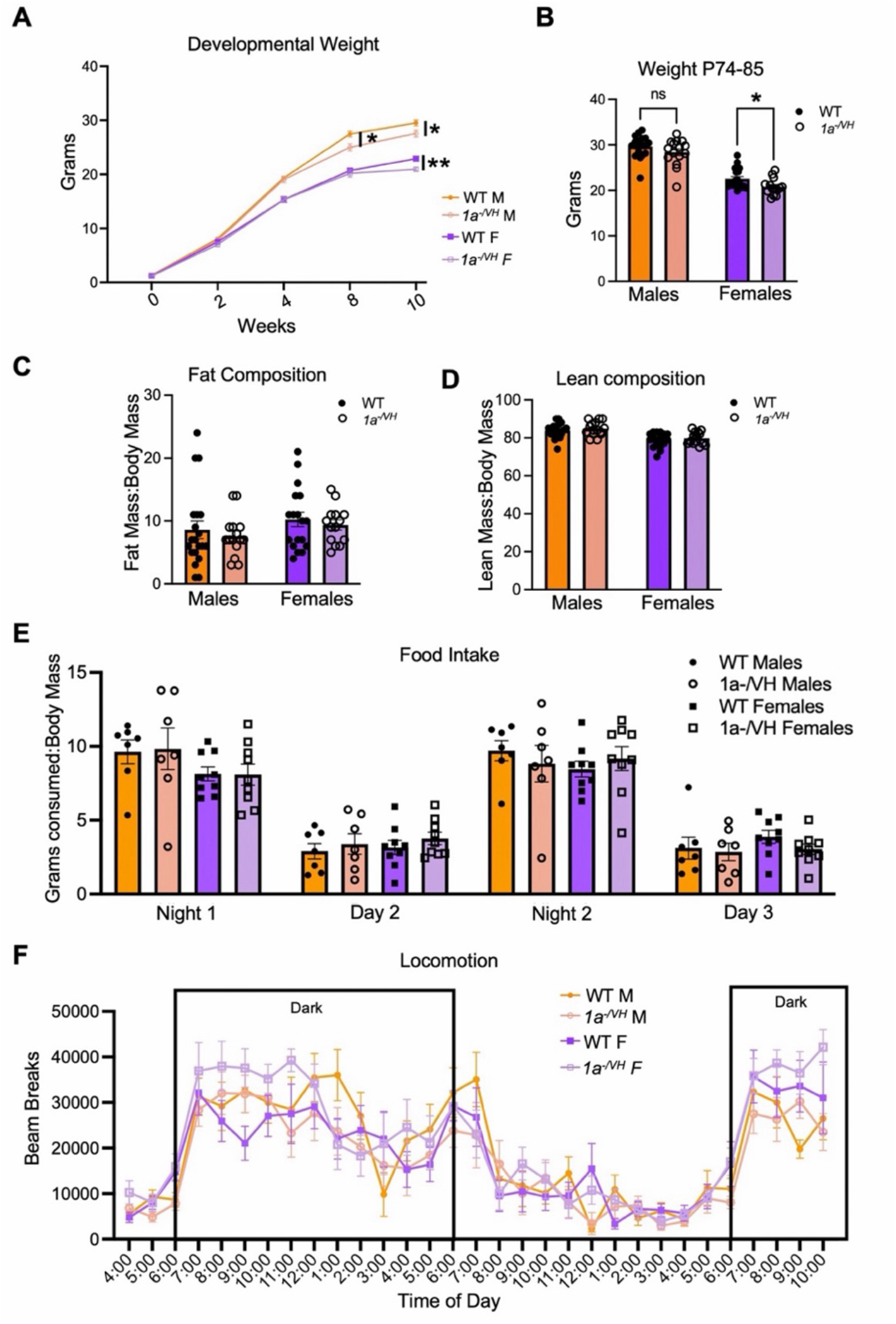
*Cc2d1a^-/VH^* mice growth deficits are not linked to body composition, or feeding. **(A)** weighing mice at different developmental timepoints showed *Cc2d1a^-/VH^* males weigh significantly less than WT littermates starting at 8 weeks old and *Cc2d1a^-/VH^* females weigh significantly less starting at 10 weeks old. **(B)** Weights of 11 week old MRI cohort mice show *Cc2d1a^-/VH^* females weight significantly less than WT littermates P74-85, while fat composition **(C)** and, lean mass composition **(D)**, are unchanged between genotype and sex. **(E)** There are no differences in food intake normalized to body mass in a metabolic chamber test. **(F)** Metabolic chamber locomotion revealed trends of increased activity in *Cc2d1a^-/VH^* females at the beginning of each dark cycle. Data are displayed as mean ± SEM. **(A-D)** Two-Way ANOVA with Fisher’s LSD. **(E-F)** Two-way ANOVA with Tukey’s multiple comparison test. \**p*<0.05, \*\**p*<0.01, \*\*\**p*<0.001. WT, wild type; P, postnatal day; ns, nonsignificant.

To investigate changes in body composition, we performed MRI-based body composition analyses to quantify fat and lean tissue mass. Male and female animals were assessed at both 9 and 11 weeks of age underscoring the subtleness of the phenotype with males showing significantly reduced weight at 9 weeks, but not 11 weeks, and females showing the opposite pattern. (**Fig. 6B**, WT M: 29.66±0.58g n=18, *1a^-/VH^* M: 28.38±0.76g n=15, *p*=0.1317, WT F: 22.54±0.52g n=18, *1a^-/VH^* F: 20.78±0.48 n=14, *p*=0.0432, **Supp. Fig 5A**). When comparing fat mass to total body mass and lean mass to total body mass we saw no differences in body composition between genotypes at either age. (**Fig. 6C-D**, Fat Composition: WT M: 7.06±1.04% n=18, *1a^-/VH^* M: 7.60±0.84% n=15, *p*=0.7034, WT F: 10.22±1.17% n=18, *1a^-/VH^* F: 9.36±0.80% n=14, *p*=0.5532, Lean composition: WT M: 83.89±0.93% n=18, *1a^-/VH^* M: 84.80±0.95% n=15, *p*=0.4839, WT F: 78.83±0.88% n=18, *1a^-/VH^* F: 79.79±0.88% n=14, *p*=0.4728, **Supp. Fig 5B-C**). These findings suggest that either the reduced body weight reflects an overall smaller body size or a more subtle biological disruption driving the weight phenotype.

Home cage feeding and motor behavior analysis was performed by housing the animals in metabolic chambers for 3 days. Comparing food consumed to body mass revealed no differences in normalized food consumption between sexes or genotypes (**Fig. 6E**, **Supp. Table 9**). Female *Cc2d1a^-/VH^* mice showed a general trend toward hyperactivity at the beginning of each dark cycle (**Fig. 6F**). Considering the mildness of this effect and the lack of body composition differences, it is unclear whether this activity difference is a key driver of the *Cc2d1a^-/VH^*female weight deficit.

We performed additional tests to assess feeding behavior in *Cc2d1a^-/VH^*animals. We first measured food consumption over a 24-hour period by providing a set amount of chow in the home cage of single-housed male and female animals. We found no genotype- or sex-dependent differences in total food intake (**Supp. Fig. 6A**). Similarly, no differences were observed in food preference between standard chow and a more palatable high-calorie, high-fat, high-sucrose chow (**Supp. Fig. 6B**).

### Distinct hypothalamic and stress signaling changes in male and female Cc2d1a hypomorph mice

Further seeking to understand whether weight was driven by metabolic changes we examined the arcuate nucleus of the hypothalamus, a key regulator of energy balance (47–49). Using qPCR, we quantified the relative expression of mRNAs for four major neuropeptides involved in appetite regulation: neuropeptide Y (NPY), agouti-related peptide (AgRP), pro-opiomelanocortin (POMC), and cocaine- and amphetamine-regulated transcript (CART). NPY and AgRP are co-expressed in orexigenic neurons that promote food intake, whereas POMC and CART are expressed in anorexigenic neurons activated by leptin signaling to promote satiety (49–53). Unexpectedly, *Cc2d1a^-/VH^*males, but not females, exhibited reduced expression of *Npy* and *Agrp* transcripts, while *Pomc* and *Cartpt* expression remained unchanged. (**Fig. 7**, *Agrp*. WT M: 1.697±0.554 n=8, *1a^-/VH^*M: 0.144±0.084 n=6, *p*=0.0039, WT F: 0.433±0.153 n=7, *1a^-/VH^*F: 0.174±0.046 n=6, *p*=0.6086, *Npy*. WT M: 1.139±0.197 n=8, *1a^-/VH^* M: 0.313±0.086 n=6, *p*=0.0182, WT F: 0.563±0.151 n=7, *1a^-/VH^* F: 0.858±0.223 n=7, *p*=0.6549, *Cartpt*. WT M: 0.943±0.114 n=6, *1a^-/VH^* M: 0.714±0.183 n=6, *p*=0.6755, WT F: 1.221±0.616 n=7, *1a^-/VH^* F: 0.988±0.177 n=6, *p*=0.6578, *Pomc*. WT M: 1.539±0.560 n=8, *1a^-/VH^* M: 0.598±0.219 n=7, *p*=0.0710, WT F: 0.547±0.219 n=7, *1a^-/VH^* F: 0.666±0.199 n=7, *p*=0.8186). These peptides play an important role in not just food intake but also nutrient partitioning and metabolic rate regulation (54–56), so it is possible *Cc2d1a^-/VH^* males have dysregulated metabolism contributing to their variable weight deficit.

**Figure 7.**
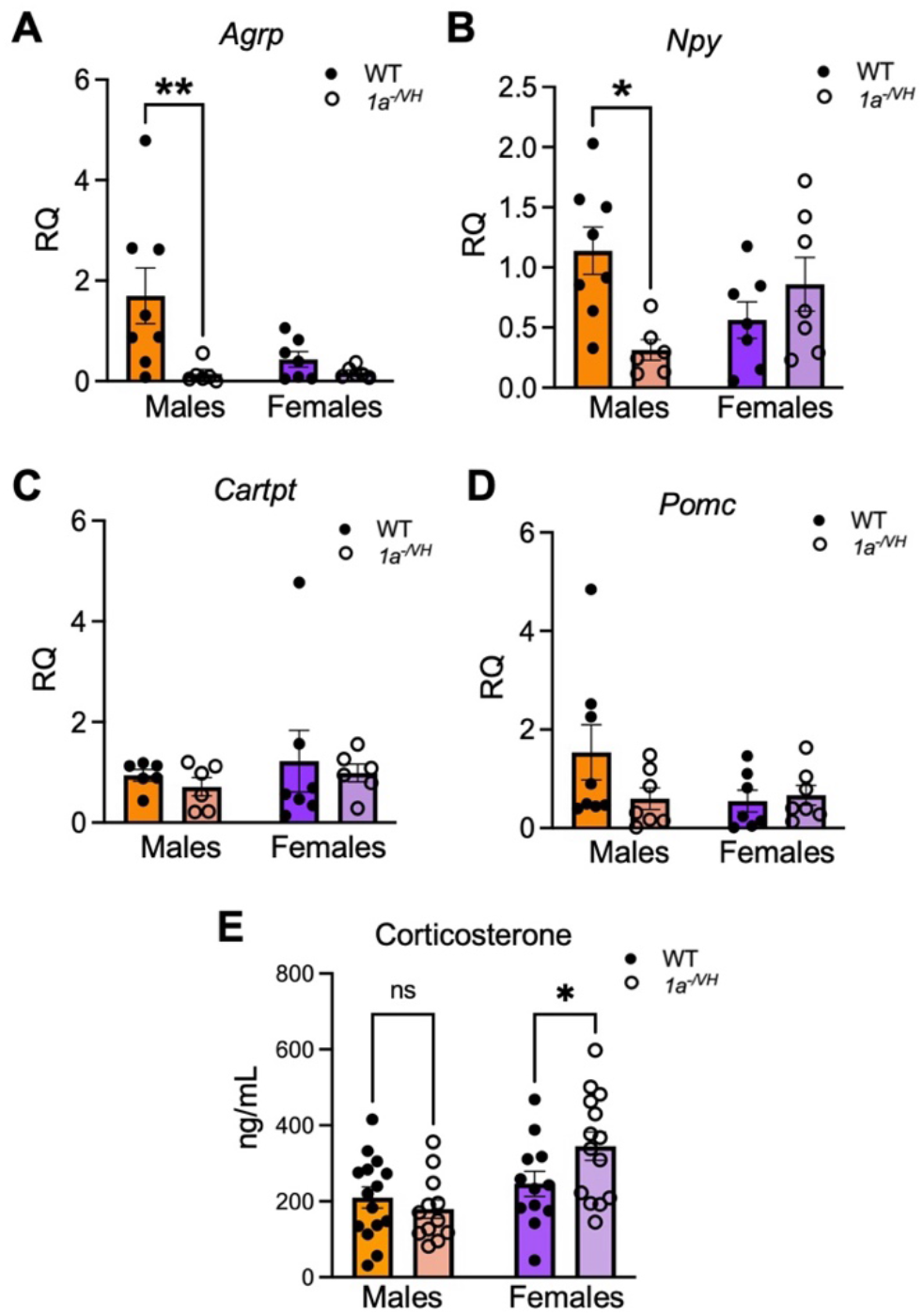
Sex-specific arcuate nucleus and corticosterone signaling in *Cc2d1a^-/VH^* mice. qPCR showed *Cc2d1a^-/VH^* males have decreased RNA transcript levels of *Agrp* **(A)** and *Npy* **(B)**. While RNA transcript levels of *Cartpt* **(C)** and *Pomc* **(D)** were unchanged between genotype and sex. **(E)** *Cc2d1a^-/VH^* females had significantly elevated serum corticosterone levels. Data are displayed as mean ± SEM. **(A-E)** Two-way ANOVA with Fisher’s LSD. WT, wild type. RQ, relative quantification normalized to WT males.

Since both AgRP and NPY can be changed in response to stress, we assessed serum corticosterone, a major physiological stress hormone that can regulate weight (57–61). In contrast, we found that *Cc2d1a^-/VH^* females showed significantly higher corticosterone levels than WT littermates, while males showed no significant difference (**Fig. 7E**, WT M: 209.81±27.76ng/mL n=15, *1a^-/VH^* M: 179.34±24.65ng/mL n=12, *p*=0.4932, WT F: 246.16±32.92ng/mL n=12, *1a^-/VH^* F: 344.87±37.14ng/mL n=14, *p*=0.0324). Chronic stress tests in rodents have been found to increase cortisol and reduce body weight (62, 63), so it is possible stress-induced weight loss may be contributing to the mutant female weight deficits.

Seeking to better understand metabolism and stress in these animals we next investigated endocrine signaling mediated by the pituitary gland, a central regulator of growth, metabolism, reproduction, and stress responses (57, 64). Using a multiplex bead-based assay, we measured circulating serum levels of several major pituitary hormones, including adrenocorticotropic hormone (ACTH), follicle-stimulating hormone (FSH), growth hormone, prolactin, and thyroid-stimulating hormone (TSH), all of which can regulate weight in sex-specific manners (57, 65–69). While there was a trend of decreased growth hormone in *Cc2d1a^-/VH^* males compared to WT littermates, no significant geno-type-dependent differences were detected for any hormone measured (**Supp. Fig 7A-J, Supp. table 10**). While these findings exclude major pituitary signaling abnormalities as the primary driver of the weight deficits, body weight regulation is highly multifactorial, and it remains possible that subtle pituitary dysfunction interacts with other biological systems to contribute to this phenotype (65, 69).

## Discussion

To our knowledge, the *Cc2d1a^VH/VH^* and *Cc2d1a^-/VH^* lines represent the first global mouse strains with profound developmental reduction of *Cc2d1a* capable of bypassing early postnatal lethality. Although promoting protein degradation was not our intent when designing the *Cc2d1a^VH/VH^*line, C-terminal HA tagging has been reported to destabilize proteins, potentially due to the increased hydrophobicity recruiting ubiquitin ligases resulting in LOF phenotypes (70–74). This serendipitous effect allowed us to determine how graded loss of *Cc2d1a* leads to cognitive and social deficits. The absence of behavioral deficits in *Cc2d1a^VH/VH^* mice reveals a striking degree of haploinsufficiency shielding, where just 20% residual CC2D1A protein is not only sufficient to circumvent the early postnatal lethality of KOs, but capable to preserve function across all behavioral measures examined. This finding supports the essential and redundant role of the CC2D1 proteins in supporting development as an allelic series in *Cc2d1a*-*Cc2d1b* double KO showed that loss of both genes leads to an empty yolk sac by embryonic day (E) 9.5 while just one allele of one of the genes is sufficient for normal embryonic development (31). The early developmental loss may allow cells to develop compensatory mechanisms that sustain cellular function such as compensatory upregulation of paralogs (75, 76). However, we detected no change in total CC2D1B levels. CC2D1B is expressed at substantially lower levels than CC2D1A and while it acts redundantly to support embryo morphogenesis in mice and flies, it cannot support postnatal survival in global *Cc2d1a* KOs (26, 31).

A plausible mechanism is homeostatic buffering, whereby cells and circuits retune their molecular operations to accommodate the loss of a specific protein. Homeostatic buffering does not restore the lost molecular function directly; rather, it stabilizes cellular output by engaging alternative molecular systems (77, 78). A well-characterized example comes from *Pax6* haploinsufficient mice, in which total excitatory synapse number remained unchanged while synapse subtypes were dynamically adjusted throughout development to preserve function (78). Given that CC2D1A loss in our lines occurs early in development, homeostatic buffering represents a compelling explanation for the lack of behavioral and molecular changes in *Cc2d1a^VH/VH^* mice. Cells may have progressively recalibrated intracellular signaling to accommodate chronically reduced CC2D1A activity and 20% residual CC2D1A may be sufficient to maintain intracellular signaling at wildtype levels.

The emergence of behavioral and intracellular signaling deficits in the 90% reduction, *Cc2d1a^-/VH^* line suggests that at this level, either the minimum threshold for sustained intracellular signaling has been crossed or homeostatic mechanisms are unable to accommodate for the molecular insult. Nevertheless, the fact that *Cc2d1a^-/VH^*mice survive and exhibit only moderate behavioral deficits, rather than postnatal lethality, indicates that 10% CC2D1A expression is sufficient to support normal respiratory function.

The behavioral profile of *Cc2d1a^-/VH^* mice recapitulates the male-biased social deficits and spatial memory acquisition delays observed in cKO models, reinforcing the importance of CC2D1A in cognitive and social function. Both CC2D1A and CC2D1B are present in synaptic fractions, but only CC2D1A is highly enriched at post-synaptic densities indicating that it may be the main paralog regulating postsynaptic signaling (32). It is plausible that the 10% threshold will primarily affect synaptic function. This is consistent with the more severe phenotypes found in *Cc2d1a*-*Cc2d1b* double heterozygote animals when compared to *Cc2d1b* KOs (31).

Postnatal forebrain *Cc2d1a* cKO lines tend to show broader behavioral disruption, including locomotor and avoidance abnormalities. In *Cc2d1a^CaMKII-Cre^* cKO mice, deficits in CREB activation were mechanistically linked to impaired spatial memory acquisition (17). By contrast, adult *Cc2d1a^-/VH^*mice showed no avoidance phenotypes or baseline CREB activation deficits *in vivo*. It is difficult to disentangle whether these additional deficits seen in cKOs reflect direct consequences of *Cc2d1a* loss or compensatory responses to its acute, cell-type-restricted removal later in development. CC2D1A is a multifunctional scaffold that interacts with many intracellular signaling pathways in different cellular compartments and cell types (1, 3, 16, 17, 22, 30, 32, 35–38). Our analysis only defines gross baseline disruptions in cortical CaMKII activation disruption in *Cc2d1a^-/VH^* males and does not address the dynamics of signaling responses. For example, studies on mouse embryonic fibroblasts obtained from *Cc2d1a* KO animals show deficits in CREB activation upon forskolin stimulation, despite equal initial baseline activity (37). What remains consistent is the male-specificity of these signaling changes supporting the role of CC2D1A in sex-specific signaling regulation.

Both CaMKII and PKA, which showed male-specific changes in *Cc2d1a* cKOs, have a well-documented interaction with sex hormone-induced plasticity. 17β-estradiol (E2) is an established modulator of hippocampal synaptic potentiation with PKA being required for initiation in females and working in coordination with CaMKII in males (79–81). Loss of CC2D1A may be disrupting this balance in males, while females with intact PKA signaling remain behaviorally unaffected. Another unexpected phenotype that could be sex hormone mediated is the mild weight deficit that emerges shortly after *Cc2d1a^-/VH^*mice reach sexual maturity (82). It is intriguing that mutant males show reduced levels of key energy balance transcripts, AgRP and NPY, in the arcuate nucleus. Transcriptional regulation of these peptides is regulated by a variety of intracellular signaling pathways triggered by hunger and satiety hormones (83). Given CC2D1A’s role as an intracellular signaling scaffold regulation it is possible that there are sex-specific intracellular signaling disruptions leading to decreased AgRP and NPY transcription. While we did not note major changes in body composition or feeding, these peptides play important roles in the regulation of nutrient partitioning and energy use that could be contributing to our subtle weight phenotype (54–56, 83, 84).

Mutant females did not show the same changes in the arcuate nucleus and there may be a different cause for their weight deficit. We found that *Cc2d1a^-/VH^* females had increased serum corticosterone levels, which could be an indicator of chronic stress. This sex-specific pattern is notable given that females generally show elevated corticosterone relative to males, and that in humans, autistic females have been reported to have higher baseline cortisol levels than autistic males (18, 85, 86). Additionally chronic stress which can be indicated by high corticosterone can cause reduced body weight in rodents (62, 63). Thus, it is possible that the reduced body weight seen in *Cc2d1a^-/VH^*females is due to chronic stress induced weight loss. This stress deficit could be subtle enough to leave body composition and feeding over all unchanged.

## Limitations

It is important to acknowledge that despite their survival these *Cc2d1a* mouse models they do not fully recapitulate the total LOF in human patients and the remaining CC2D1A expression may be causing compensatory effects. While our studies suggest a more severe effect of *CC2D1A* LOF on male cognition and sociability and an effect on weight gain, there is no clinical data available in humans comparing males and females with CC2D1A variants or investigating other symptoms beyond an ID and ASD diagnosis. There is only one report of a case with a homozygous *CC2D1A* variant in a study of syndromic short stature (87).

Overall, the scope of this study was to determine a threshold for CC2D1A expression and to confirm whether males were still preferentially affected with developmental protein reduction. We were not powered to detect deficits with small effect size and to statistically confirm sex-specificity, which will require up to 4 times the current sample size. In experiments with the traditional sample size used for behavioral analyses (10-15 animals), the mild weight deficits were difficult to assess consistently, and we could not compare males and females directly. Larger Ns will be needed in studies focused on weight and sex-specificity. Further work will be needed to determine the mechanisms behind CaMKII modulation and the possible disruption of AgRP and NPY in reduced male weight.

## Conclusions

In summary, this study introduces two novel *Cc2d1a* hypomorph strains and reveals a strikingly narrow expression threshold governing the penetrance of *Cc2d1a* LOF phenotypes. The preservation of normal behavior at 20% residual protein demonstrates that even partial restoration of CC2D1A levels may yield meaningful therapeutic benefit. The *Cc2d1a^-/VH^* line further establishes that loss of CC2D1A has more deleterious effects on male intracellular signaling and social and cognitive behavior. The puberty onset weight deficits with reduced arcuate nucleus peptide transcript levels in mutant males, but not females, together raise the intriguing possibility that CC2D1A could also play a sex-specific role in growth and metabolism. Beyond modeling *CC2D1A LOF* pathology with greater developmental and cellular fidelity, *Cc2d1a^-/VH^* mice offer a novel platform for investigating the molecular mechanisms of sex-specific phenotypes in neuropsychiatric illnesses.

## Data Availability

All numerical data presented in the manuscript and raw western blot images are available on Zenodo at doi: 10.5281/zenodo.23087973.

## Abbreviations

ACTH: Adrenocorticotropic hormone
AgRP: Agouti-related peptide
ASD: Autism Spectrum Disorder
CaMKII: Calcium/calmodulin-dependent protein kinase II
CART: Cocaine and amphetamine-regulated transcript
CC2D1A: Coiled-coil and C2 Domain Containing 1A
CC2D1B: Coiled-coil and C2 Domain Containing 1B
cKO: Conditional knock out
CREB: Cyclic AMP response element-binding protein
ERK1/2: Extracellular Signal-Regulated Kinase 1 and 2
FSH: Follicle stimulating hormone
ID: Intellectual disability
KO: Knock out
MRI: Magnetitic resonance imaging
MWM: Morris Water Maze
NPY: Neuropeptide Y
PDE4D: Phosphodiesterase 4D
PKA: Protein kinase A
POMC: Pro-opiomelanocortin
TSH: Thyroid Stimulating hormone
V5/HA: Simian Virus 5/hemagglutinin
WT: Wild Type

## Author contributions

ATH, AB, and MCM conceptualized the study and defined the methodology; ATH, AB, PR, SR, VU, ES, and SW performed experiments and collected data; ATH, AB, and MCM analyzed the data; ATH and MCM visualized the data, drafted and edited the original manuscript; MCM acquired funding for the study; all authors reviewed the final draft of the manuscript.

## Acknowledgements

We would like to thank all the members of the Manzini laboratory for their insights during the course of this project. We would like to thank Drs. Mark Rossi, Ben Samuels, Gleb Shumyatsky and Jim Millonig for discussions on metabolism, behavior, and modeling neuropsychiatric disorders. We are grateful to Dr. Christoph Buettner for access to the EchoMRI Body Composition Analyzer and to Dr. Jim Millonig for access to the Luminex 200 system for Milliplex analysis.

