## Supplementary data for "Dose dependent impact of global *Cc2d1a* reduction on cognitive, social, and metabolic phenotypes"

Heller et al.

**Supplementary Figure 1.**

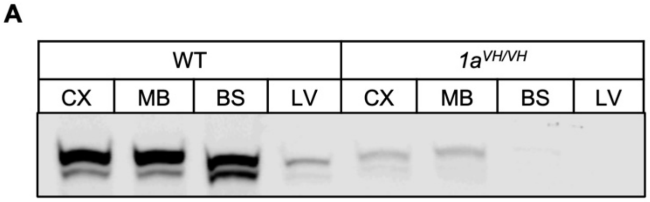


**Supplementary Figure 1. Homozygous insertion of the V5-HA tag (*Cc2d1a*$^{VH/VH}$) leads to global protein reduction.** Representative western blot image probed for CC2D1A in both WT and *Cc2d1a*$^{VH/VH}$ mice across different brain regions (CX, cortex; MB, midbrain; BS, brainstem) and the liver (LV) show global CC2D1A reduction.

**Supplementary Figure 2.**

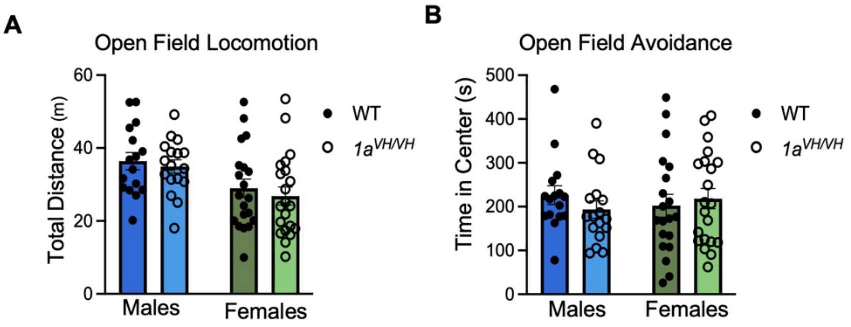


**Supplementary Figure 2. *Cc2d1a*$^{VH/VH}$ mice do not display deficits in the open field test.** *Cc2d1a*$^{VH/VH}$ mice show comparable **(A)** total distance traveled, locomotion, and **(B)** avoidance of center time in the open field test. Data are displayed as mean ± SEM. Two-way ANOVA with Fisher's LSD. WT, Wild type

**Supplementary Figure 3.**

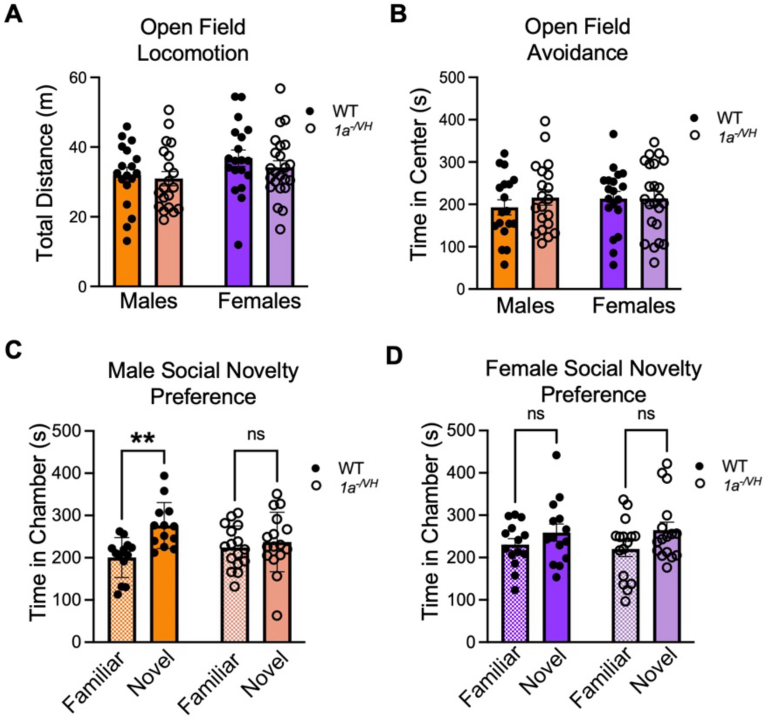


**Supplementary Figure 3. *Cc2d1a$^{-/VH}$* mice have social novelty, but not open field deficits.** **(A)** *Cc2d1a$^{-/VH}$* mice show comparable **(A)** total distance traveled, locomotion, and **(B)** avoidance of center time in the open field test. **(C)** *Cc2d1a$^{VH/VH}$* males, but not females **(D)** show decreased novel mouse interaction time in 3-chamber social-novelty preference testing. Data are displayed as mean ± SEM. Two-way ANOVA with Fisher's LSD. **$p<0.01$. WT, wild type.

**Supplementary Figure 4.**

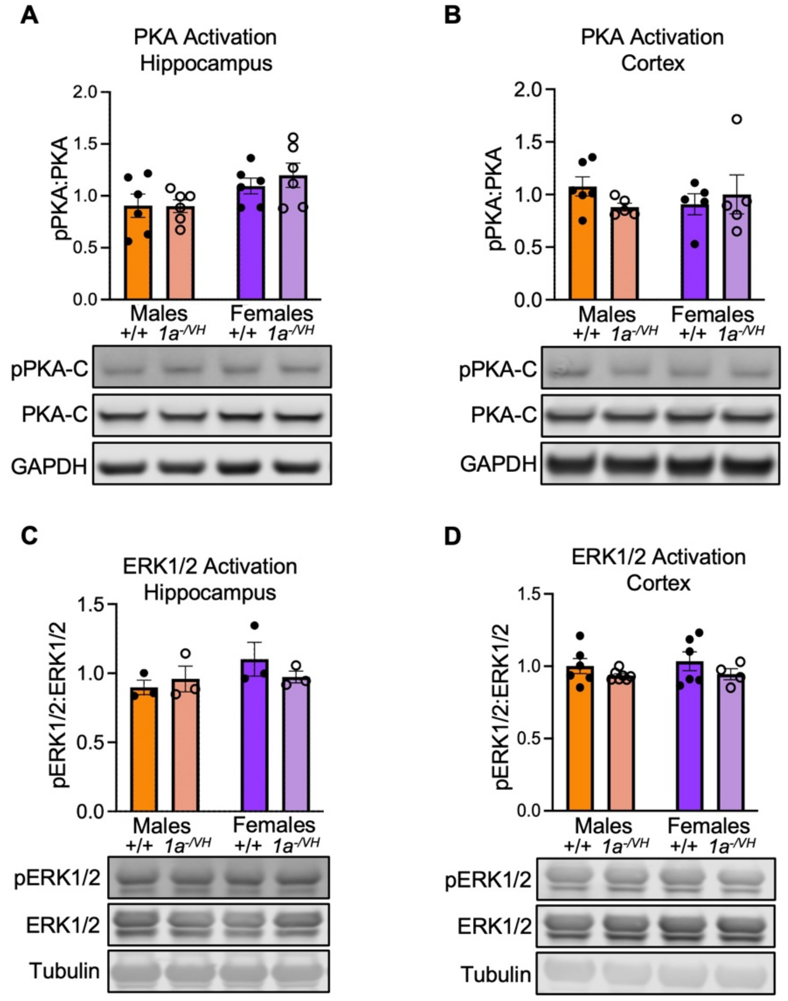


**Supplementary Figure 4. *Cc2d1a*$^{-/VH}$ mice do not have baseline PKA or ERK activation deficits.** Western blot shows *Cc2d1a*$^{-/VH}$ mice have no differences in baseline PKA activation in either adult hippocampus **(A),** or cortex **(B)**. Western blot shows *Cc2d1a*$^{-/VH}$ mice have no differences in baseline ERK1/2 activation in either adult hippocampus **(C),** or cortex **(D)**. Data are displayed as mean ± SEM. Two-way ANOVA with Fisher's LSD. WT, Wild type pPKA-C, T197 phosphorylated PKA-C; pERK1/2, T202 phosphorylated ERK1 Y204 phosphorylated ERK2.

**Supplementary Figure 5.**

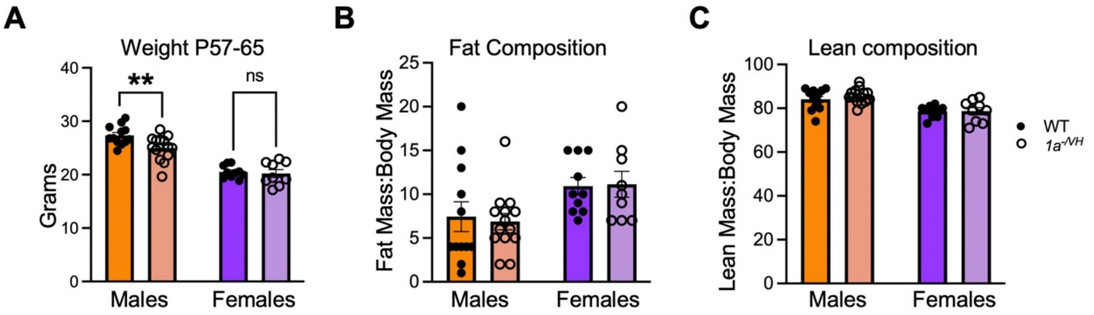


**Supplementary Figure 5. *Cc2d1a$^{-/VH}$* mice do not show body composition differences at 9 weeks old. (A)** Weights of 9 week old MRI cohort mice show that *Cc2d1a$^{-/VH}$* males weigh significantly less than WT littermates P57-65 while fat composition **(B)** and, lean mass composition **(C)**, are unchanged between genotype and sex. Data are displayed as mean ± SEM. Two-way ANOVA with Fisher's LSD. WT, Wild type

**Supplementary Figure 6.**

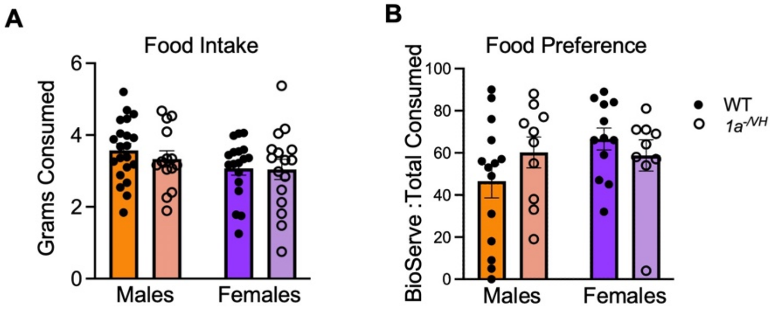


**Supplementary Figure 6. *Cc2d1a$^{-/VH}$* mice do not have home cage feeding deficits. (A)** There are no differences in 24 hour total food intake between genotype and sex. **(B)** Nor are there differences in food 24 hour food preference between standard 5058 chow and BioServe Supreme Mini-Treats. Data are displayed as mean ± SEM. Two-way ANOVA with Fisher's LSD. WT, Wild type

### Supplementary Figure 7.

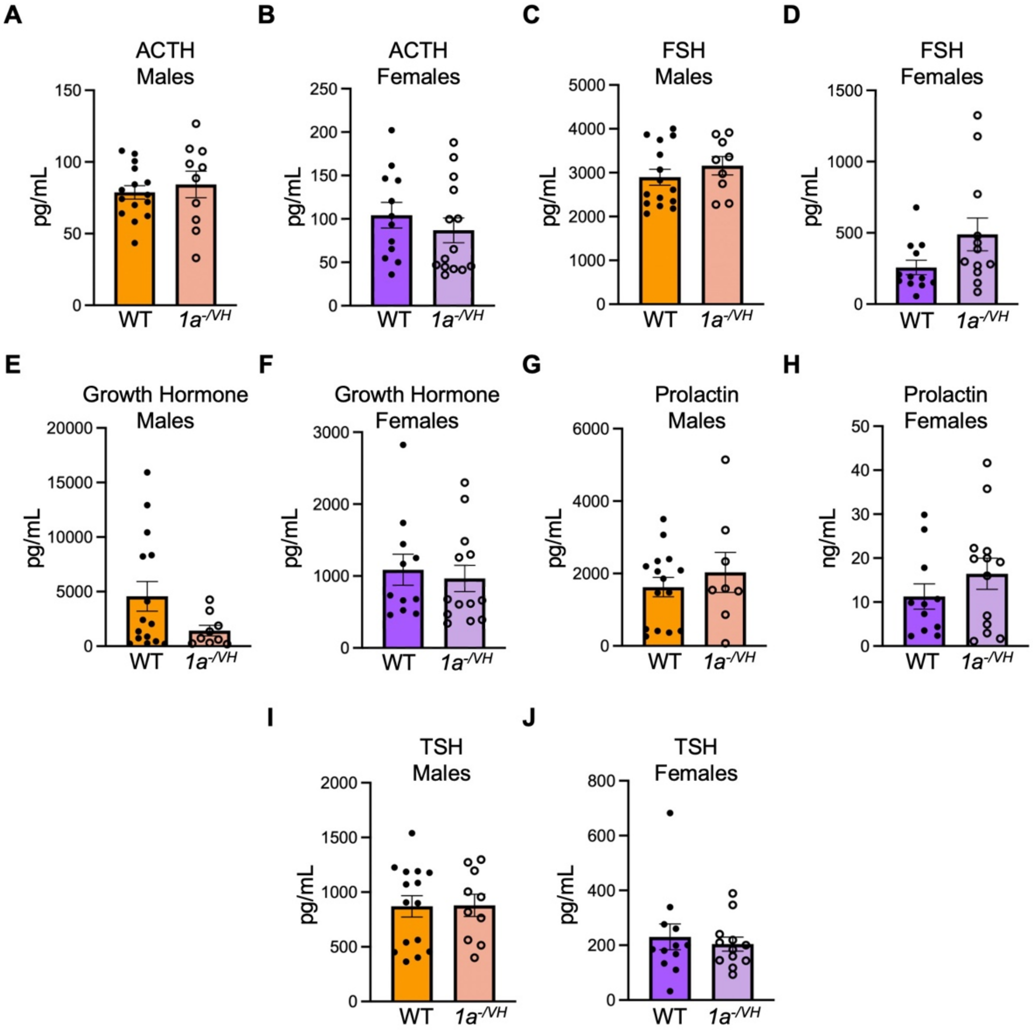


**Supplementary Figure 7. *Cc2d1a*$^{-/VH}$ mice have normal pituitary gland signaling.** There are no differences between genotype in blood serum hormone levels of ACTH in males **(A)** or females **(B)**, FSH males **(C)** or females **(D)**, growth hormone males **(E)** or females (F), prolactin males **(G)** or females **(H)** and TSH in males **(I)** or females **(J)**. Data are displayed as mean ± SEM. Mann-Whitney U test. WT, wild type; ACTH, adrenocorticotropic hormone; FSH, follicle stimulating hormone; TSH, thyroid stimulating hormone.

**Supplementary Table 1.** *Cc2d1a*$^{VH/VH}$ Line SHIRPA Characterization

| Genotype | Days postnatal | Weight (g) | Rightin g Reflex Present [a] | Reaching Reflex Present [b] | Wire Hang (s) | Tail Pinch Present [c] | Ear Reflex Present [d] | Whisker Reflex Present [d] | Eye Reflex Present [d] |
|---|---|---|---|---|---|---|---|---|---|
| WT M | 89.58±3.00 | 30.42±0.59 | 12/12 | 12/12 | 24.58±7.07 | 12/12 | 12/12 | 10/12 | 10/12 |
| *1a*$^{VH/VH}$ M | 93.00±3.78 | 29.07±0.91 | 11/11 | 11/11 | 25.23±7.07 | 11/11 | 11/11 | 10/11 | 10/11 |
| WT F | 90.25±2.24 | 23.85±0.60 | 12/12 | 12/12 | 35.73±7.44 | 12/12 | 12/12 | 11/12 | 11/12 |
| *1a*$^{VH/VH}$ F | 93.94±1.93 | 24.45±0.87 | 18/18 | 18/18 | 38.84±5.86 | 17/18 | 18/18 | 16/18 | 16/18 |

[a] fraction of animals that righted themselves. [b] fraction of animals that had the visual reaching reflex. [c] fraction of animals that responded to pain pinch. [d] fraction of animals that responded to auditory stimulus

**Supplementary Table 2.** *Cc2d1a*$^{VH/VH}$ 3-chamber sociability and social novelty

| Stage | Genotype | Interaction Type | Mean±SEM (s) | n | *p*-value |
|---|---|---|---|---|---|
| Sociability | WT M | Empty | 202.36±16.70 | 10 | 0.0075 |
| Sociability | | Mouse | 295.38±22.29 | | |
| Sociability | *1a*$^{VH/VH}$ M | Empty | 191.69±12.82 | 11 | 0.0030 |
| Sociability | | Mouse | 292.34±11.45 | | |
| Sociability | WT F | Empty | 186.02±9.14 | 19 | <0.0001 |
| Sociability | | Mouse | 291.98±9.72 | | |
| Sociability | *1a*$^{VH/VH}$ F | Empty | 189.53±10.52 | 18 | 0.0002 |
| Sociability | | Mouse | 271.67±15.99 | | |

**Supplementary Table 3.** *Cc2d1a*$^{VH/VH}$ Morris Water Maze

| Trial | **WT M (s)** n=9 | ***1a*$^{VH/VH}$ M (s)** n=12 | **Male *p*-value** | **WT F (s)** n=12 | ***1a*$^{VH/VH}$ F (s)** n=10 | **Female *p*-value** |
|---|---|---|---|---|---|---|
| H1 | 18.46±4.11 | 18.56±2.00 | >0.9999 | 18.66±4.10 | 24.70±2.53 | 0.9743 |
| H2 | 12.84±3.60 | 14.48±2.55 | >0.9999 | 15.24±2.27 | 23.37±6.32 | 0.9256 |
| H3 | 13.23±2.84 | 12.54±2.48 | >0.9999 | 12.28±1.39 | 11.77±1.68 | >0.9999 |
| H4 | 8.53±1.32 | 16.18±3.71 | 0.4946 | 14.82±2.50 | 13.79±3.39 | >0.9999 |
| H5 | 9.06±1.17 | 10.91±2.12 | 09958 | 10.34±1.72 | 13.26±2.19 | 0.9638 |
| H6 | 7.86±1.27 | 8.82±1.33 | 0.9998 | 10.55±1.60 | 10.74±1.62 | >0.9999 |
| H7 | 8.58±1.03 | 9.94±1.69 | 0.9981 | 7.45±1.21 | 8.39±0.84 | 0.9989 |
| H8 | 6.84±1.32 | 7.04±0.91 | >0.9999 | 8.72±1.58 | 7.92±0.95 | >0.9999 |
| H9 | 5.74±0.59 | 6.58±0.96 | 0.9966 | 7.55±1.28 | 5.97±0.60 | 0.9491 |
| Probe[a] | 30.02±1.86 | 30.21±2.87 | 0.9598 | 32.44±1.81 | 30.32±3.22 | 0.5549 |
| R1 | 44.91±9.40 | 55.85±7.96 | 0.8586 | 67.67±8.79 | 64.00±8.07 | 0.9968 |
| R2 | 14.82±3.28 | 27.80±8.87 | 0.5730 | 31.64±8.63 | 16.22±4.49 | 0.4327 |
| R3 | 8.56±1.99 | 13.83±3.98 | 0.6895 | 13.72±2.89 | 9.36±2.86 | 0.7562 |
| R4 | 8.96±2.05 | 10.43±3.55 | 0.9942 | 8.13±1.90 | 14.51±5.88 | 0.7918 |

Time (s) to escape platform. [a] seconds spent in escape quadrant.

**Supplementary Table 4.** *Cc2d1a*$^{-/VH}$ Line SHIRPA Characterization

| Genotype | Days postnatal | Weight (g) | Righting Reflex Present [a] | Reaching Reflex Present [b] | Wire Hang (s) | Tail Pinch Present [c] | Ear Reflex Present [d] | Whisker Reflex Present [d] | Eye Reflex Present [d] |
|---|---|---|---|---|---|---|---|---|---|
| WT M | 72.78±0.46 | 29.67±0.70 | 9/9 | 9/9 | 17.09±5.70 | 7/9 | 9/9 | 7/9 | 7/9 |
| *1a*$^{-/VH}$ M | 73.56±0.18 | 27.67±0.97 | 9/9 | 9/9 | 34.60±8.60 | 9/9 | 9/9 | 9/9 | 9/9 |
| WT F | 78.47±2.26 | 23.96±0.51 | 15/15 | 15/15 | 47.66±4.76 | 15/15 | 14/15 | 14/15 | 14/15 |
| *1a*$^{-/VH}$ F | 79.94±2.29 | 21.46±0.28 | 18/18 | 18/18 | 46.53±5.34 | 17/18 | 18/18 | 18/18 | 18/18 |

[a] fraction of animals that righted themselves. [b] fraction of animals that had the visual reaching reflex. [c] fraction of animals that responded to pain pinch. [d] fraction of animals that responded to auditory stimulus.

**Supplementary Table 5.** *Cc2d1a$^{-/VH}$* 3-chamber sociability and social novelty statistical results

| Stage | Genotype | Interaction Type | Mean±SEM (s) | n | *p*-value |
|---|---|---|---|---|---|
| Sociability | WT M | Empty | 186.49±14.90 | 13 | 0.0026 |
| Sociability | | Mouse | 313.68±17.79 | | |
| Sociability | *1a$^{-/VH}$* M | Empty | 217.05±18.42 | 16 | 0.1323 |
| Sociability | | Mouse | 270.66±21.39 | | |
| Sociability | WT F | Empty | 180.66±11.22 | 14 | <0.0001 |
| Sociability | | Mouse | 334.53±12.93 | | |
| Sociability | *1a$^{-/VH}$* F | Empty | 190.80±12.01 | 17 | <0.0001 |
| Sociability | | Mouse | 318.93±12.39 | | |
| Social Novelty | WT M | Familiar Mouse | 200.43±13.08 | 13 | 0.0153 |
| Social Novelty | | Novel Mouse | 276.28±15.06 | | |
| Social Novelty | *1a$^{-/VH}$* M | Familiar Mouse | 224.43±12.73 | 16 | 0.6279 |
| Social Novelty | | Novel Mouse | 237.38±17.58 | | |
| Social Novelty | WT F | Familiar Mouse | 230.67±14.25 | 14 | 0.4386 |
| Social Novelty | | Novel Mouse | 258.96±20.32 | | |
| Social Novelty | *1a$^{-/VH}$* F | Familiar Mouse | 220.36±17.91 | 16 | 0.1976 |
| Social Novelty | | Novel Mouse | 264.79±18.79 | | |

**Supplementary Table 6.** *Cc2d1a$^{-/VH}$* Morris Water Maze statistical results

| Trial | **WT M (s)** n=15 | ***1a$^{-/VH}$* M (s)** n=17 | **Male *p*-value** | **WT F (s)** n=13 | ***1a$^{-/VH}$* F (s)** n=13 | **Female *p*-value** |
|---|---|---|---|---|---|---|
| H1 | 16.67±1.97 | 20.64±2.95 | 0.6671 | 22.47±4.06 | 24.56±6.04 | >0.9999 |
| H2 | 13.16±2.02 | 17.38±2.40 | 0.5896 | 15.12±2.40 | 12.98±3.90 | >0.9999 |
| H3 | 10.29±2.02 | 15.25±1.77 | 0.3677 | 13.85±2.51 | 12.36±1.92 | >0.9999 |
| H4 | 12.72±1.96 | 13.67±1.33 | 0.9996 | 14.01±2.23 | 11.50±1.72 | 0.9868 |
| H5 | 9.97±1.45 | 13.10±2.36 | 0.8857 | 8.71±0.99 | 10.88±1.42 | 0.8960 |
| H6 | 9.43±0.95 | 14.16±2.16 | 0.4340 | 9.62±2.00 | 9.53±1.41 | >0.9999 |
| H7 | 10.91±1.31 | 10.58±1.52 | >0.9999 | 7.64±1.14 | 7.68±0.85 | >0.9999 |
| H8 | 9.98±1.45 | 8.72±1.00 | 0.9998 | 7.37±1.07 | 7.78±1.17 | >0.9999 |
| H9 | 7.21±0.65 | 8.39±0.85 | 0.9999 | 6.92±0.57 | 7.24±0.91 | >0.9999 |
| Probe[a] | 26.62±1.61 | 24.53±2.00 | 0.4110 | 25.86±1.74 | 24.51±1.99 | 0.6290 |
| R1 | 57.75±7.31 | 53.36±6.22 | 0.9851 | 41.06±7.66 | 53.39±7.83 | 0.7180 |
| R2 | 22.50±4.00 | 32.18±4.94 | 0.4493 | 25.50±6.64 | 28.79±6.77 | 0.9948 |
| R3 | 12.77±2.60 | 22.34±5.42 | 0.4145 | 22.12±6.53 | 23.66±7.27 | 0.9998 |
| R4 | 7.97±1.19 | 17.36±5.13 | 0.3202 | 18.13±6.36 | 14.68±3.03 | 0.9814 |

**Supplementary Table 7.** *Cc2d1a$^{-/VH}$* Morris Water Maze Last Trial of Day ANOVA results

| Trial | **WT M (s)** n=15 | ***1a$^{-/VH}$* M (s)** n=17 | **Male *p*-value** | **WT F (s)** n=13 | ***1a$^{-/VH}$* F (s)** n=13 | **Female *p*-value** |
|---|---|---|---|---|---|---|
| H3 | 10.29±2.02 | 15.25±1.77 | 0.0509 | 13.85±2.51 | 12.36±1.92 | 0.9545 |
| H6 | 9.43±0.95 | 14.16±2.16 | 0.0675 | 9.62±2.00 | 9.53±1.41 | >0.9999 |
| H9 | 7.21±0.65 | 8.39±0.85 | 0.9180 | 6.92±0.57 | 7.24±0.91 | 0.9881 |

Time (s) to escape platform. [a] seconds spent in escape quadrant.

**Supplemental Table 8.** *Cc2d1a*$^{-/VH}$ Developmental Weights

| Week | WT M (g) | *1a*$^{-/VH}$ M (g) | Male *p*-value | WT F (g) | *1a*$^{-/VH}$ F (g) | Female *p*-value |
|---|---|---|---|---|---|---|
| 0 | 1.34±0.11, n=3 | 1.27±0.05, n=7 | >0.9999 | 1.23±0.06, n=5 | 1.19±0.07, n=5 | >0.9999 |
| 2 | 8.08±0.22, n=9 | 7.72±0.35, n=9 | 0.9979 | 7.51±0.22, n=12 | 7.02±0.40, n=12 | 09516 |
| 4 | 19.34±0.30, n=10 | 19.04±0.61, n=9 | 0.9990 | 15.34±0.57, n=9 | 15.29±0.48, n=12 | >0.9999 |
| 8 | 27.47±0.55, n=11 | 24.98±0.67, n=13 | 0.0182 | 20.71±0.36, n=9 | 20.21±0.72, n=9 | 0.9718 |
| 10 | 29.51±0.54, n=21 | 27.55±0.66, n=19 | 0.0152 | 22.89±0.46, n=22 | 20.95±0.33, n=18 | 0.0011 |

**Supplemental Table 9.** Home cage feeding statistical results

| | **WT M** n=7 | ***$1a^{-/VH}$* M** n=7 | **Male *p*-value** | **WT F** n=9 | ***$1a^{-/VH}$* F** n=9 | **Female *p*-value** |
|---|---|---|---|---|---|---|
| Night 1 | 9.64±0.80 | 9.84±1.41 | 0.9993 | 8.14±0.47 | 8.10±0.71 | >0.9999 |
| Day 2 | 2.91±0.53 | 3.39±0.70 | 0.9442 | 3.16±0.49 | 3.77±0.42 | 0.7755 |
| Night 2 | 9.71±0.68 | 8.83±1.24 | 0.9238 | 8.46±0.53 | 9.17±0.80 | 0.8795 |
| Day 3 | 3.11±0.74 | 2.86±0.59 | 0.9931 | 3.88±0.44 | 3.04±0.36 | 0.4716 |

Food intake : body mass

**Supplemental Table 10.** Pituitary gland hormone quantification

| | **WT M (pg/mL)** | ***$1a^{-/VH}$* M (pg/mL)** | **Male *p*-value** | **WT F (pg/mL)** | ***$1a^{-/VH}$* F (pg/mL)** | **Female *p*-value** |
|---|---|---|---|---|---|---|
| ACTH | 78.76±4.74, n=15 | 84.29±9.28, n=10 | 0.5305 | 104.3±14.84, n=12 | 86.9±14.25, n=14 | 0.2740 |
| FSH | 2896±180.6, n=15 | 3160±209.5, n=9 | 0.3786 | 257.2±50.47, n=12 | 489.7±114.8 n=12 | 0.0780 |
| GH | 4567±1352, n=15 | 1425±485.3, n=9 | 0.2152 | 1087±216.6, n=11 | 965.3±182.8, n=13 | 0.4940 |
| Prolactin | 1625±269.6, n=15 | 2031±551.9, n=8 | 0.7284 | 11281±2876, n=11 | 16441±3543, n=13 | 0.4585 |
| TSH | 870.3±97.20, n=15 | 878.5±101.8, n=10 | 0.8490 | 230.5±47.05, n=12 | 204.1±25.44, n=12 | 0.9774 |
